# Cross-Frequency Multilayer Network Analysis with Bispectrum-based Functional Connectivity: A Study of Alzheimer’s Disease

**DOI:** 10.1101/2021.08.07.455499

**Authors:** Dominik Klepl, Fei He, Min Wu, Daniel J. Blackburn, Ptolemaios G. Sarrigiannis

## Abstract

Alzheimer’s disease (AD) is a neurodegenerative disorder known to affect functional connectivity (FC) across many brain regions. Linear FC measures have been applied to study the differences in AD by splitting neurophysiological signals, such as electroencephalography (EEG) recordings, into discrete frequency bands and analysing them in isolation from each other. We address this limitation by quantifying cross-frequency FC in addition to the traditional within-band approach. Cross-bispectrum, a higher-order spectral analysis approach, is used to measure the nonlinear FC and is compared with the cross-spectrum, which only measures the linear FC within bands. This work reports the reconstruction of a cross-frequency FC network where each frequency band is treated as a layer in a multilayer network with both inter- and intra-layer edges. Cross-bispectrum detects cross-frequency differences, mainly increased FC in AD cases in *δ*-*θ* coupling. Overall, increased strength of low-frequency coupling and decreased level of high-frequency coupling is observed in AD cases in comparison to healthy controls (HC). We demonstrate that a graph-theoretic analysis of cross-frequency brain networks is crucial to obtain a more detailed insight into their structure and function. Vulnerability analysis reveals that the integration and segregation properties of networks are enabled by different frequency couplings in AD networks compared to HCs. Finally, we use the reconstructed networks for classification. The extra cross-frequency coupling information can improve the classification performance significantly, suggesting an important role of cross-frequency FC. The results highlight the importance of studying nonlinearity and including cross-frequency FC in characterising AD.

## I. INTRODUCTION

Alzheimer’s disease (AD), the most common form of dementia, causes early degradation of neural circuits leading to cell death and synaptic loss [1]. Studies have shown that AD affects distributed brain networks and alters functional connectivity (FC), which can lead to disconnection syndrome and disrupts information processing across multiple scales [2–5].

Electroencephalography (EEG) is a common method to study AD. Currently, EEG is not commonly used for diagnosing AD, but many studies demonstrate high potential in developing EEG-based biomarkers and diagnostic tools. The main EEG characteristics associated with AD are the slowing of signals, and altered synchronisation [3, 5–8]. Slowing of EEG in AD was observed as increased power in *δ* and *θ* frequency bands and decreased power in *α* and *β* frequency bands [5, 6]. Similarly, AD shows increased synchronisation within low-frequency bands (*<*12 Hz), decreased synchronisation in high-frequency bands, and is associated with the altered FC, especially the long-distance cortical connections [9]. However, these characteristics are typically measured at a single channel or between channel pairs. In contrast, network-based methods analyse multiple channels and reveal additional characteristics of AD, namely, reduced integration of information [10, 11], and loss of small-worldness [12]. However, these characteristics are often analysed only within specific frequency bands.

This study aims to extend the FC beyond within-frequency coupling (WFC), taking the cross-frequency coupling (CFC) [13] into account. WFC networks of AD were analysed previously by using coherence (linear) [14] and wavelet coherence (nonlinear) [15]. Only one CFC measure, i.e. phase synchronisation index (PSI), had been used for the graph analysis of CFC networks in AD [16]. This work extended the findings of reduced integration and loss of small-worldness to CFC multilayer networks. However, it does not consider the roles of different frequency components in the networks. The multilayer-network framework had been used previously for brain network analysis. Loss of inter-frequency hubs in AD was reported using MEG multiplex networks[17, 18], where the inter-layer edges are inserted with fixed weight only between the same nodes across layers. Alterations in multilayer network hubs have been reported in multi-modal networks in AD [19], and fMRI frequency-band networks in schizophrenia [20]. Multilayer networks integrating WFC and CFC have been used to analyse MEG data from healthy [21], and schizophrenic subjects [22]. Tewarie et al. [21] show that frequency-band network layers interact via CFC, share a certain amount of structure and operate at the edge of independence and interdepen-dence. However, these studies analyse the layer relationships mainly as the correlation of their adjacency matrices or as differences in global average coupling strength.

Bispectrum is a higher-order spectral analysis and quantifies quadratic coupling between two frequency components and their algebraic sum [23]. It has been shown to detect amplitude-amplitude and phase-amplitude CFC in addition to the phase-phase coupling [24, 25]. The bispectral coupling also indicates an increase in non-Gaussianity [26]. Features derived from bis-pectrum were proposed as biomarkers of epilepsy [27, 28], Parkinson’s disease [29], autism [30] and AD [26, 31, 32]. Most of these studies compute (cross-) bispectra of only a few channels or pairs of channels. Although a few studies used bispectrum to compute global networks from multiple channels [33], these analyses do not use graph theory. Instead, each node is analysed in isolation [31] or single-channel bispectra are averaged across nodes to derive certain global properties [32]. In contrast, this study computes cross-bispectra between all pairs of EEG channels to estimate the widely distributed FC brain networks and perform graph-theoretical analysis.

In this work, the cross-bispectrum (CBS) estimates of FC are computed. We aim to investigate the contribution of nonlinear WFC and CFC in differentiating between Alzheimer’s disease (AD) and healthy controls (HC) in comparison to the equivalent linear WFC measured with cross-spectrum (CS) (Fig. 1). We report a multilayer network-based analysis to elucidate the roles of the traditional EEG frequency bands and their CFC in the sensor-level EEG networks of HC and AD. Moreover, we use the reconstructed brain networks to classify AD using a support vector machine classifier.

**FIG. 1:**
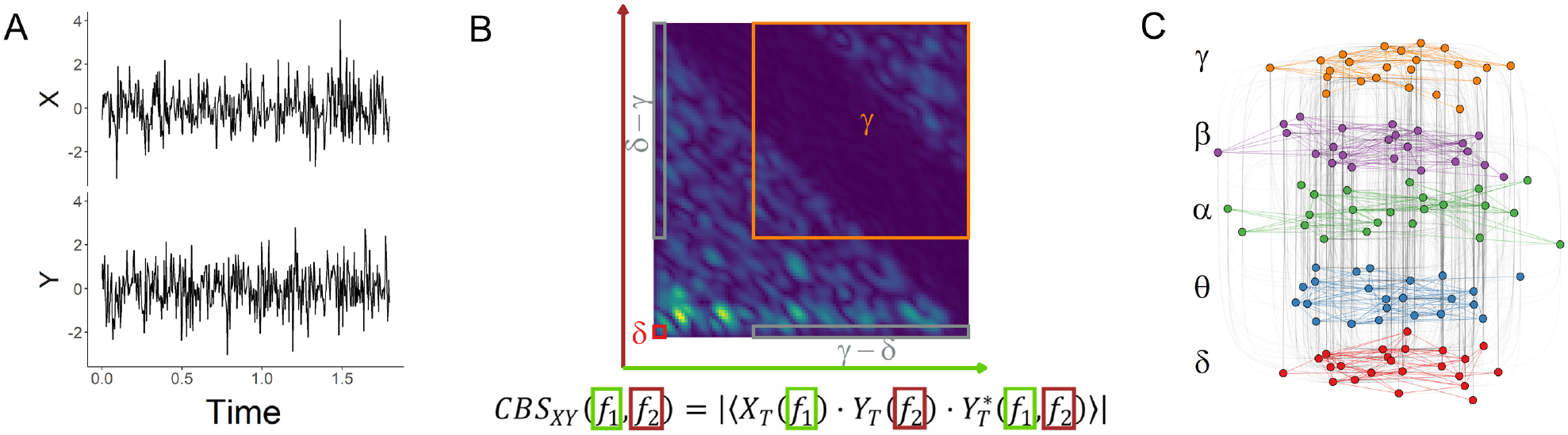
A conceptual schematic of implementing the proposed cross-bispectrum (CBS) multilayer network analysis. (A) Each EEG signal is cleaned and scaled. (B) For each pair of EEG electrodes, a cross-bispectrum is estimated. The frequency bands coupling edge weights are given by the average value within the respective CBS window, e.g. *δ*-*δ* (red). Note that CBS estimates are directed, e.g. *δ*-*γ ̸*= *γ*-*δ* (both in grey). Thus from each CBS, 25 edges are inferred. (C) Using the edge weights inferred from CBS, a multilayer network is constructed with layers representing the frequency bands of EEG. Such a network has both intra-layer and inter-layer edges, representing within-frequency and cross-frequency coupling, respectively.

## II. DATA

EEG recordings were collected from 20 AD patients and 20 healthy control participants (HC) younger than 70 years. A detailed description of the experimental design and confirmation of the diagnosis is provided in [34]. All AD participants were recruited in the Sheffield Teaching Hospital memory clinic. AD participants were diagnosed between 1 month and 2 years before data collection, and all of them were in the mild to moderate stage of the disease at the time of recording with the average Mini Mental State Examination score of 20.1 (sd = 4). Age and gender-matched HC participants with normal neuropsychological tests and structural MRI scans were recruited.

EEG was acquired using an XLTEK 128-channel head-box, Ag/AgCL electrodes with a sampling frequency of 2 kHz using a modified 10-10 overlapping a 10-20 international electrode placement system with a referential montage with a linked earlobe reference. The recordings lasted 30 minutes, during which the participants were instructed to rest and not to think about anything specific. Within each recording, there were two-minute-long epochs during which the participants had their eyes closed (alternating with equal duration eyes-open epochs, not used in this work).

All the recordings were reviewed by an experienced neurophysiologist on the XLTEK review station with time-locked video recordings (Optima Medical LTD) to isolate artefact-free segments. For each participant, three 12-second-long artefact-free epochs were isolated. Finally, the following 23 bipolar channels were created: F8–F4, F7–F3, F4–C4, F3–C3, F4–FZ, FZ–CZ, F3–FZ, T4–C4, T3–C3, C4–CZ, C3–CZ, CZ–PZ, C4–P4, C3–P3, T4–T6, T3–T5, P4–PZ, P3–PZ, T6–O2, T5–O1, P4–O2, P3–O1 and O1–O2 [34].

### A. EEG pre-processing

EEG signals were confirmed to be artifact-free. Thus, no additional artifact removal was undertaken. The signals were band-pass filtered to be between 0.1 and 100 Hz using a zero-phase 5^th^ order Butterworth filter; 50 Hz relating to the power line noise was removed using a zero-phase 4^th^ order Butterworth stop-band filter, and the data were down sampled to 250 Hz using an order 8 Chebyshev type I filter. Finally, the signals were normalised (to zero mean and unit standard deviation).

## III. METHODS

### A. Cross-spectrum and cross-bispectrum

The spectrum *S_X_* of a signal *X* is calculated via smoothed periodogram. Fast Fourier Transform (FFT) is used to estimate the periodogram with Daniell smoothers. The periodogram is computed over 256 frequency bins (0.98 Hz bandwidth). CS at frequency *f* is then computed as: *CS_XY_* (*f*) = *S_X_* (*f*) · *S_Y_* (*f*). An absolute value of CS is calculated. A direct FFT-based method is used to estimate the absolute value of CBS:

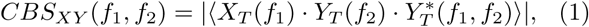

where ⟨·⟩ denotes averaging, *X_T_* (*f*) is an FFT of signal *X* over an interval *T* and *Y ^∗^* is the complex conjugate. 256-point FFT is used. CBS is computed over 1-second-long windows with 50% overlap over the whole frequency range (0.5 - 100 Hz). The window size and overlap were chosen empirically to balance the spectral and temporal resolutions. The estimated CBS is then smoothed in the frequency domain using the Rao-Gabr window (size 5).

CS and CBS were computed for all pairs of EEG channels. Five frequency bands *b* are considered: *δ* (0.5 − 5*Hz*), *θ* (5 − 8*Hz*), *α* (8 − 16*Hz*), *β* (16 − 32*Hz*) and *γ* (32 − 100*Hz*).

The connectivity between channels *X* and *Y* and frequency bands *b_X_* and *b_Y_* is computed as:

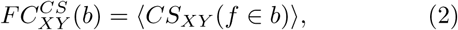

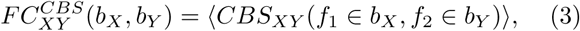

for CS and CBS, respectively, where ⟨·⟩ denotes averag-ing. This resulted in five WFC (CS and CBS) and 20 CFC (CBS only) measures per channel pair. It is of note that the CBS is directed.

In order to ensure the reliability of the estimated connectivity, surrogate thresholding was used [35]. For each pair of channels, 200 surrogate signals were generated using the FFT surrogate, which scrambles the phase of the signal, and their CS and CBS are computed. The 95% confidence interval of surrogate values is computed and used as a threshold. Coupling values below the threshold are set to zero. We chose this approach to ensure the reliability of estimated brain networks. In contrast, Wang et al. [31] used no such thresholding when analysing bi-coherence coupling. Alternative approaches exist in the literature. Chella et al. [33] take advantage of the asymmetric property of CBS to ensure robustness against mixing artefacts.

We obtain a set of connectivity matrices for each EEG recording, i.e. *N* × *N* matrices (*N* = 23). For CS and CBS, there are five and 25 connectivity matrices, respectively. A global (averaged per subject) connectivity is computed for each 23 × 23 matrix and compared between groups using a two-sample t-test if normally distributed and a Mann-Whitney test otherwise.

### B. Network measures

To identify the important channels in the network, we compute a coupling-specific node strength [36] for each channel *i* and different types of frequency couplings *c*,

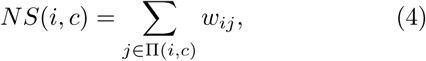

where Π(*i, c*) are the nodes connected to channel *i* via edge type *c* and *w_ij_* is the edge weight, i.e. CS or CBS connectivity given by *ij*th entry of the *N* ×*N* connectivity matrix. This measure is computed for both CS and CBS, resulting in 5 (5 frequency bands) and 25 (5×5 frequency bands) values per channel, respectively.

In order to analyse the importance of the different frequency couplings within the global brain network, we represent them as a multilayer network. In this network, nodes are located within layers representing the different frequency bands. WFC represents the edges between nodes within a single layer, i.e. intra-layer, and CFC represents the edges between nodes located in different layers, i.e. inter-layer. In this paper, the CS networks are not analysed as multilayer networks since such networks would have no inter-layer edges and thus would not be comparable directly with the CBS networks. The following measures are computed only for CBS networks. We obtain networks with 23 nodes that are replicated over 5 layers (*L* ∈ [*δ, θ, α, β, γ*]), resulting effectively in 115 nodes. There are 5 types of intra-layer edges, such as *δ*-*δ*, and 20 types of inter-layer edges, such as *δ*-*θ* or *θ*-*δ*.

We measure the importance of each type of frequency coupling within the multilayer network by measuring the contribution of each edge to enable the efficient passing of information through the network. For this purpose, we define coupling betweenness centrality (CBW) based on an adjusted version of edge betweenness [37]:

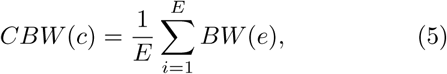

where *E* is the total number of edges of coupling type *c* and *BW* (*e*) is edge betweenness centrality given by:

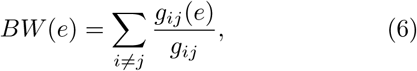

where *g_ij_* is the number of shortest paths between nodes *i* and *j*, and *g_ij_*(*e*) is the number of those paths that go through edge *e*. The shortest path is defined as a path with the least sum of 1*/w_ij_*. CBW quantifies the contribution of each coupling type to the information integration [38], i.e. the amount of information flow through edges. Note that the CBW of a weighted and unweighted version of the same network results in different value of CBW. Therefore, we analyse both weighted and un-weighted CBW.

CBW assumes that the essential processes within the network occur along the shortest paths. However, there might be alternative paths with only minor length differences, which CBW ignores. In case of a disruption of the network structure, these alternative paths might enable the recovery of function with negligible differences. We quantify this as the vulnerability of the network to the removal of one type of frequency coupling. The vulnerability is measured in two ways: the loss of ability to integrate information [39] and the loss of segregation.

The integration property of network *G*, i.e. the ability of a network to communicate information globally, is approximated with global efficiency (*E_G_*) given by:

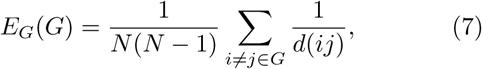

where *N* is the number of nodes in network *G* and *d*(*ij*) is the shortest path length between nodes *i* and *j*. *E_G_*is related to CBW. CBW measures the information flow on the more detailed edge level while *E_G_* takes the node-level perspective.

The segregation property of network *G*, i.e. the presence of densely connected clusters and sparse connections between them, is approximated with a local efficiency given by:

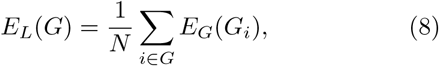

where *G_i_* is the neighbourhood of node *i*, i.e. subgraph of nodes directly connected to *i*, without node *i* itself.

In order to measure the vulnerability of the network and its dependence on different types of frequency coupling, *E_G_* and *E_L_* are computed for the full network. The two measures are then re-computed for a perturbed network where one type of frequency coupling (i.e. a set of edges) is removed. The change in *E_G_* and *E_L_*give the global and local vulnerability measures *V_G_*(*G_c_*) = 1 − (*E_G_*(*G_c_*)*/E_G_*(*G*)) and *V_L_* = 1 − (*E_L_*(*G_c_*)*/E_L_*(*G*)), where *G* is the full network and *G_c_* is the perturbed network with the edges of coupling type *c* removed.

### C. Network thresholding and statistical analysis

In order to filter out the unimportant edges that might result from a spurious coupling, the weighted multilayer networks are thresholded through relative quantile-based thresholding. Given a quantile *Q*, all edges with a weight lower than *Q* are removed from the network. There are considerable differences between the weights of each frequency coupling type (e.g. mean of *γ*-*β* = 1.627 compared to mean of *α*-*α* = 8.975); thus, a separate threshold *Q* is used. As a result, the networks retain *Q*% of the strongest edges. To ensure that the observed differences between the networks are not due to the choice of threshold *Q*, all of the network measures are computed over 20 threshold values (*Q* ∈ [0, 0.95] in increments of 0.05), and only significant differences observed over at least ten thresholds are declared significant. The reported plots and numerical results are generated from such threshold levels of *Q* that the between-group difference is maximised (i.e. largest effect size). However, the effect of choice of this threshold should be minimal as significant differences were observed over multiple thresholds. All *p*-values are corrected using the Benjamini–Hochberg false discovery rate method [40].

Additionally, to improve the reliability, we perform epoch-wise test-retest experiments. For each participant included in this study, there are three epochs. Thus, we repeat the full analysis reported in this paper for each epoch separately. Consequently, only significant differences observed consistently across all three epochs are denoted as significant. An analysis of statistical power given our sample size was performed to identify the threshold effect size where 80% is reached (A).

Furthermore, we convert the CBS network from directed to undirected by taking the mean weight for each pair of directed edges, thus collapsing them into a single edge. Such an approach is the most conservative since the potential effect of outliers is minimised compared to the alternative of taking the maximum weight.

Node strength is log-transformed to reduce skewness. For node strength, we do not threshold the networks as this can lead to isolated nodes with no edges. We test whether node strengths are normally distributed for each coupling and channel separately with a Shapiro test. Node strengths that pass the test are then compared with a two-sample t-test, and those that do not pass the test are compared with a Mann-Whitney U test.

The multilayer graph measures such as CBW, *V_G_* and *V_L_* aim to analyse the roles of frequency coupling types in terms of the network’s properties. However, it is unclear whether such multilayer networks should be weighted or unweighted. Thus, we examine the patterns in both weighted and unweighted multilayer graphs. The weighted networks can be converted into unweighted networks by setting the weights of all edges to 1. Additionally, the selected graph metrics, except for node strength, assume edge weights represent the distance between weights. Since functional connectivity is a measure of similarity, we convert the edge weights to distance as follows,

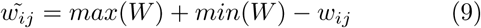

,where 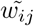 is the transformed edge weight connecting nodes *i* and *j*, *W* is the edge weight distribution of the graph and *max*(·) and *min*(·).

We test whether *CBW*, *V_G_* and *V_L_* of both weighted and unweighted networks are normally distributed using the Shapiro-Wilk test for each coupling type separately. A two-sample t-test is used for normally distributed variables and Mann-Whitney U for non-normally distributed variables to compare between groups. Furthermore, both weighted and unweighted *CBW* and *V_G_* are log-transformed to reduce skewness.

### D. Network classification

Finally, we train classifiers using the network metrics to evaluate the predictive power of these biomarkers of AD. Three classifiers are trained using the CS, CBS, and combined features, respectively. In other words, the CS classifier is trained using node strength, the CBS classifier uses the node strength and multilayer network metrics, and the combined classifier uses all of the previous. Additionally, these features are collected across all filtered networks.

As this leads to a large feature space, we introduce an effect-size-based forward feature selection. The features are ordered by the absolute value of effect size (Cohen’s *d* [41]) and sequentially added to the feature vector, which is then used to train the classifier. The first 100 features are evaluated in this manner. Note that comparing the CS and CBS classifiers is likely unfair as the CS utilises considerably smaller and less complex features, as the node strength is a relatively simple network measure. Instead, the CS classifier should be viewed as a naive baseline.

Support vector machine classifier with radial basis kernel is used as the classifier. Moreover, features are scaled to zero mean and unit standard deviation. 10-fold cross-validation repeated 100 times is used to train and evaluate the classifier.

Finally, we use the feature sets of CS and CBS classifiers that achieved the best performance and train a combined classifier. We hypothesise that the information captured by CS and CBS networks is at least partially unique. Thus a classifier trained on the combined feature sets should outperform the classifiers trained on individual networks, as it can leverage the information from both functional connectivity measures.

## IV. RESULTS AND DISCUSSION

We denote a statistical test as significant only if it is consistently detected across at least ten network thresholds and in all three epochs. Therefore, for simplicity, only results from epoch two are reported in the following sections, except for the classification results, where data from all epochs are utilised. Epoch two was selected randomly, which does not affect the reported results as all results were required to be observed across all three epochs. Moreover, for visualisation purposes, we select the network threshold, where the strongest difference is observed for each comparison separately.

The results and visualisations from epochs 1 and 3 are included in B and D, respectively. The numerical results from epoch 2 are included in C.

### A. Connectivity matrices and average connectivity

Differences in averaged connectivity matrices (Fig. 2 and 3) indicate that both methods seem to detect varia-tions in the topology of FC networks. The results of sta-tistical tests are reported in C (Tables C.XI and C.XII). By using CS, significant differences in the average con-nectivity are found in *δ* and *θ* bands, where AD cases have increased connectivity. Additionally, CS reveals a decrease in *β* connectivity of AD cases.

**FIG. 2:**
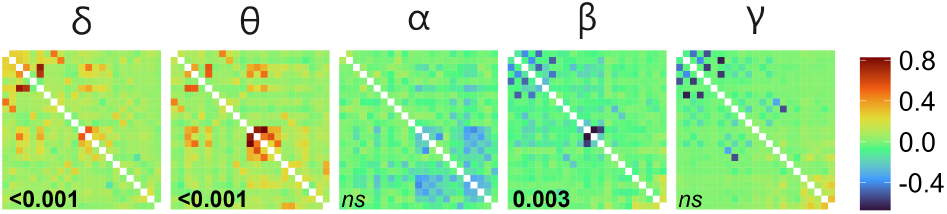
The difference between average connectivity matrices (*AD − HC*) measured with cross-spectrum in epoch 2. For visualisation purposes, the values were min-max normalised. Digits in black denote a *p*-value (FDR corrected) testing for the difference in global coupling (*p <* 0.05 in bold, in italics otherwise).

**FIG. 3:**
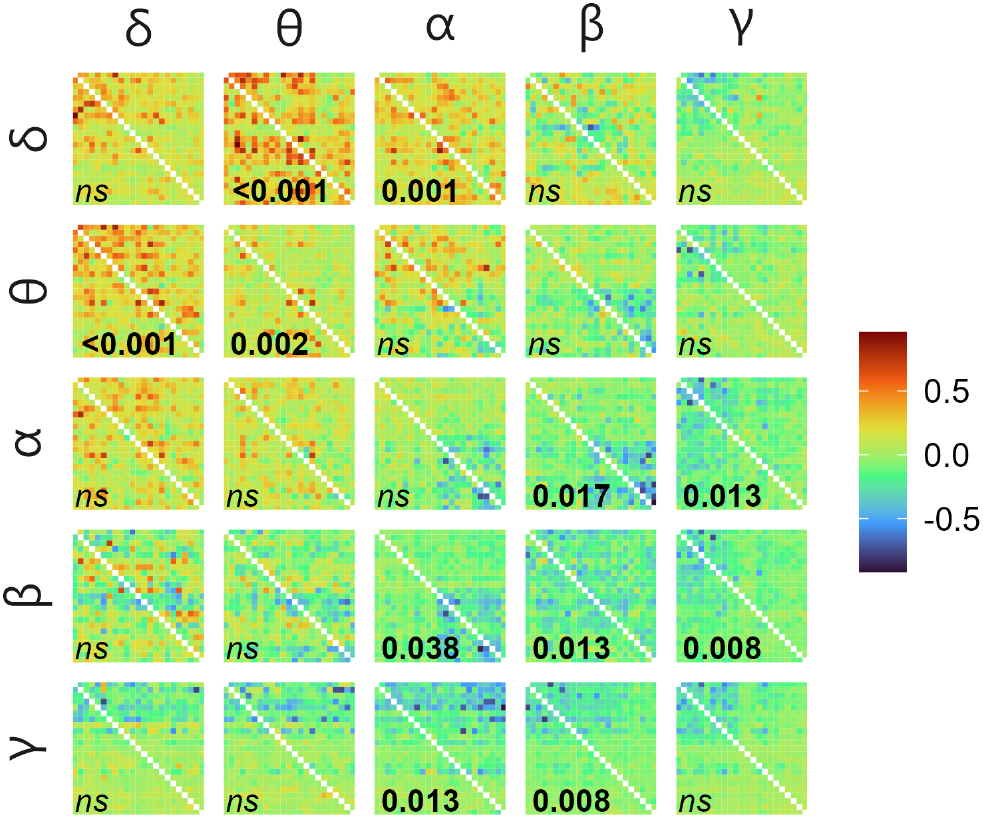
The difference between average connectivity matrices (*AD − HC*) measured with cross-bispectrum in epoch 2 with input frequency on the vertical facets and output frequency on the horizontal. For visualisation purposes, the values were min-max normalised. Digits in black denote a *p*-value (FDR corrected) testing for the difference in global coupling (*p <* 0.05 in bold, in italics otherwise).

Using CBS, differences can be observed in multiple frequency bands and their couplings. Increased global connectivity is observed in AD cases in *θ* WFC and *θ*-*δ*, *δ*-*θ* and *δ*-*α* CFC. In contrast, decreased global connectivity in AD cases is found in *β* WFC and alpha-beta, *α*-*γ*, *β*-*α*, *β*-*γ*, *γ*-*α* and *γ*-*β* CFC. Overall, AD cases show increased connectivity in low-frequency components and their CFC interactions and decreased connectivity in high-frequency components.

These findings are consistent with the literature reporting increased activity in *δ* and *θ* in AD [3, 5]. An increase in *δ* WFC and low-frequency CFC in AD was also reported using bicoherence [31]. Similarly, Maturana et al. [32] report increased bispectral power in AD in *δ* and *θ* and a decrease in *α*, *β*_1_ and *β*_2_. They also report lower bispectral entropy in *δ* and *θ* suggesting fewer frequency components interact with these frequency bands. In contrast, Cai et al. [16] report the opposite differences in the same WFC and CFC using PSI, i.e. decrease in *δ* and *θ*.

Moreover, the visible structure distortion within multiple frequency bands detected by both CS and CBS suggests connections to the disconnection syndrome and dis-turbed information processing in AD.

### B. Coupling-wise node strength

In order to statistically test the differences in connectivity measured by both CS and CBS and to localise the brain regions which show the most pronounced differences between AD and HC, node strength is measured for each channel and coupling type separately. We show the results in Figs. 2 and 3 for CS and CBS, respectively. The details of these statistical tests are reported in C (Tables C.XIII and C.XIV).

The differences in WFC detected by CS and CBS (Fig. 4 and diagonal elements in Fig. 5) are generally similar. Both methods show increased *θ* node strength in AD cases across most channels. Both CS and CBS show decreased *β* coupling. However, each detects these changes in different regions, i.e. CS across all channels except for occipital, while CBS only in central channels. Interestingly, CBS fails to capture the increased node strength in *δ* in AD cases that can be seen in CS. These differences showcase the importance of assessing both linear and nonlinear coupling in understanding the variations in AD brain networks.

**FIG. 4:**
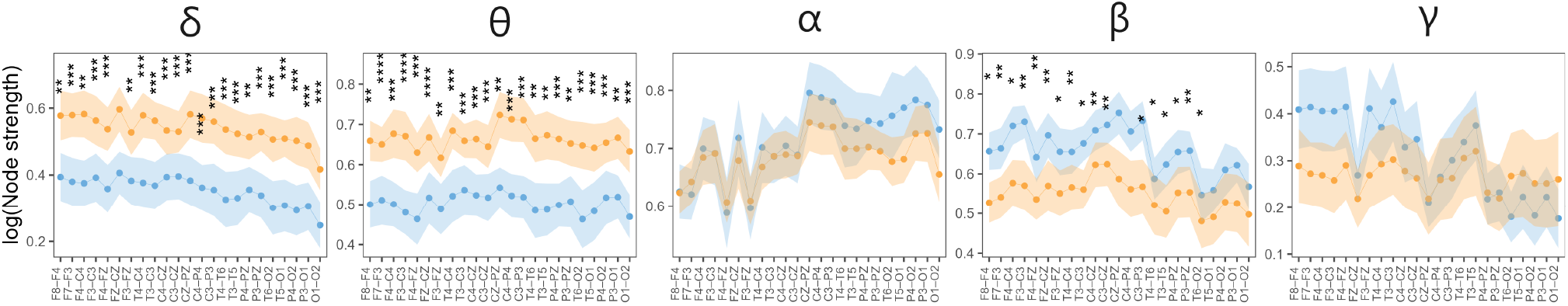
Node strength (min-max normalised) measured with CS of HC (blue) and AD (orange): mean with 95% confidence intervals. Significant differences observed in at least ten thresholded networks and across all epochs are encoded by asterisks. The number of asterisks corresponds to the p-value (FDR corrected), i.e. *p ≤* 0.0001 “****”, *p ≤* 0.001 “***”, *p ≤* 0.01 “**”, and *p ≤* 0.05 “*”.

Multiple differences in the CFC (off-diagonal elements in Fig. 5) are detected, highlighting the need to analyse the interactions of frequency components in both healthy and AD brain networks. AD cases show a global increase in *δ*-*θ* and *θ*-*δ*, and in frontal and temporal areas in *δ*-*α*.

**FIG. 5:**
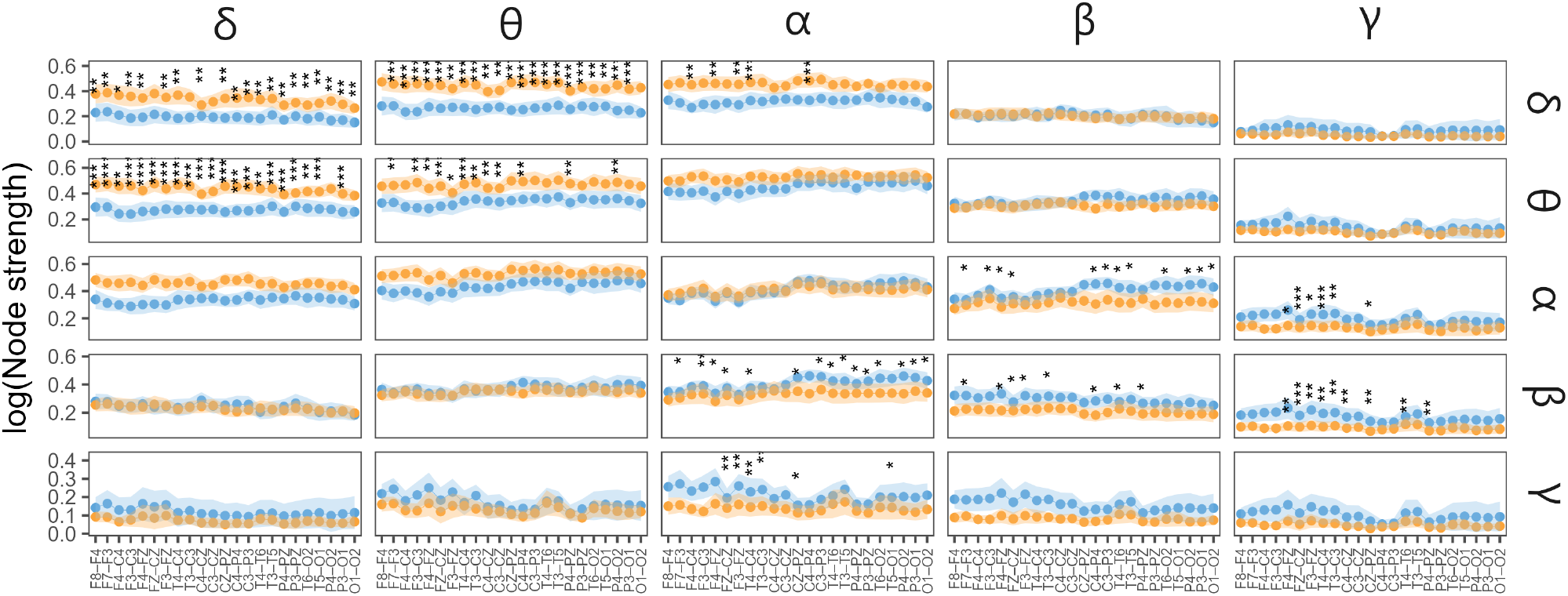
Node strength (min-max normalised) measured with CBS of HC (blue) and AD (orange): mean with 95% confidence intervals. The input frequency is on the vertical facets, and the output frequency is on the horizontal. Significant differences observed in at least ten thresholded networks and across all epochs are encoded by asterisks. The number of asterisks corresponds to the p-value (FDR corrected), i.e. *p ≤* 0.0001 “****”, *p ≤* 0.001 “***”, *p ≤* 0.01 “**”, and *p ≤* 0.05 “*”.

This is in contrast to the findings in Wang et al. [31], where an increase of *δ*-*θ* only in frontal channels is reported. They also report an increase in midline parietal-occipital *θ*-*γ* that we do not detect. Furthermore, we observe a frontal, occipital and temporal decrease in *α*-*β* and *β*-*α*, frontocentral and frontotemporal decrease in *α*-*γ* and *β*-*γ*, and in frontal, frontocentral and occipital channels in *γ*-*α* in AD cases.

Cai et al. [16] report comparable differences using PSI, but in contrast to our results, they show mainly decreased node strength in AD cases. This might be because CBS is influenced by the amplitude, while PSI is a pure phase coupling measure. Fraga et al. [42] report an increase of the *δ*-*θ* and *δ*-*β* amplitude-amplitude CFC in AD cases which is similar to our results. This suggests that CBS indeed measures some mixture of CFC types [24] since our results are partially in line both with phase-phase and amplitude-amplitude CFC studies.

### C. Multilayer network analysis

In order to elucidate the roles of the frequency bands and their coupling, both WFC and CFC, we analyse the CBS networks as multilayer networks with five layers rep-resenting the traditional frequency bands of EEG. More-over, both the weighted and unweighted versions of these networks are analysed.

First, weighted and unweighted CBW are used to as-sess the importance of each type of coupling for both local and global communications in the network. Re-sults of statistical tests comparing the unweighted CBW are reported in the appendix (Table C.XV) and visu-alised in Fig. 6. Results of statistical tests comparing the weighted CBW are reported in the appendix (Table C.XVIII) and visualised in Fig. 7.

**FIG. 6:**
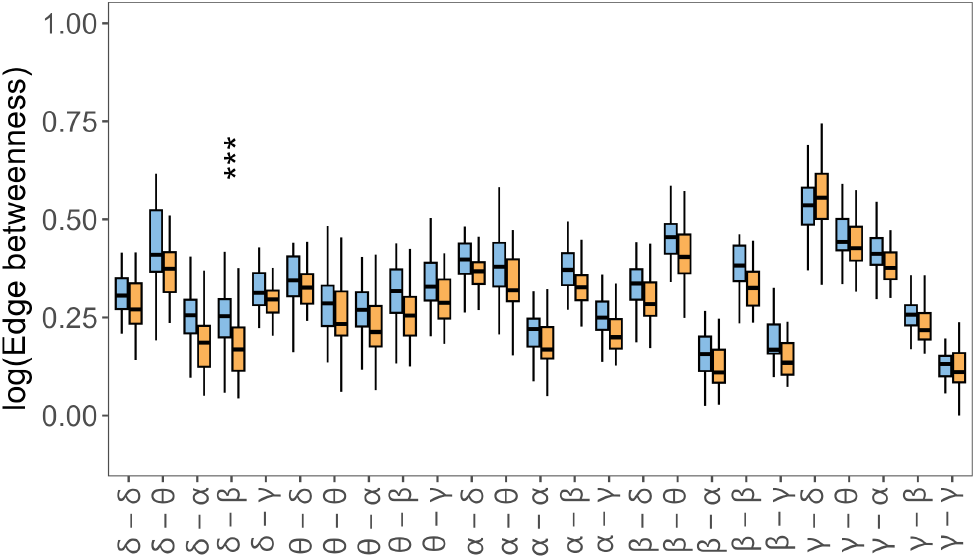
Importance of each type of frequency coupling of HC (blue) and AD (orange) measured by edge betweenness. Significant differences observed in at least ten thresholded networks and across all epochs are encoded by asterisks. The number of asterisks corresponds to the p-value (FDR corrected), i.e. *p ≤* 0.0001 “****”, *p ≤* 0.001 “***”, *p ≤* 0.01 “**”, and *p ≤* 0.05 “*”.

**FIG. 7:**
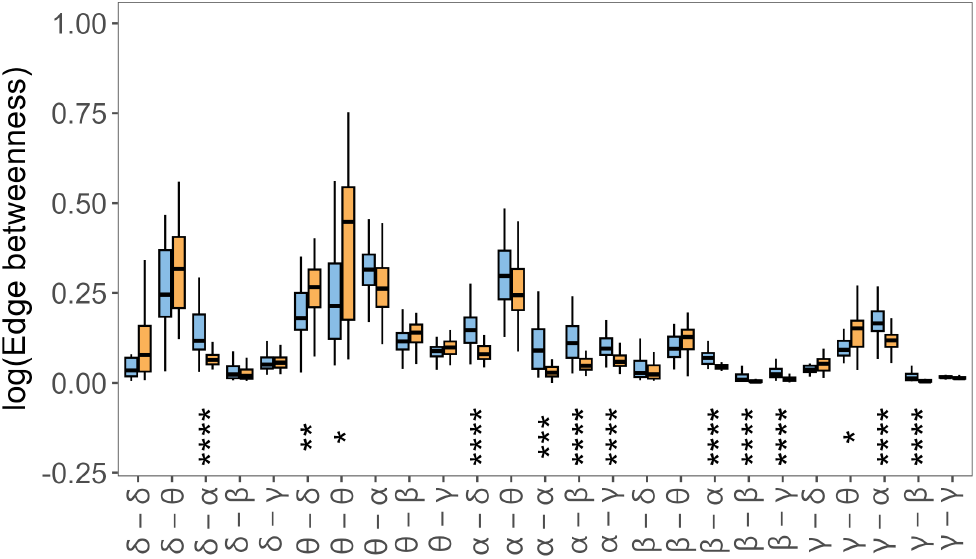
Importance of each type of frequency coupling of HC (blue) and AD (orange) measured by weighted edge betweenness. Significant differences observed in at least ten thresholded networks and across all epochs are encoded by asterisks. The number of asterisks corresponds to the p-value (FDR corrected), i.e. *p ≤* 0.0001 “****”, *p ≤* 0.001 “***”, *p ≤* 0.01 “**”, and *p ≤* 0.05 “*”.

The unweighted CBW shows only a decrease in AD cases in the *δ*-*β* CFC (Fig 6. In contrast, the weighted CBW shows multiple decreases in AD cases, specifically in *α*-*α* and *β*-*β* WFC and *δ*-*α*, *α*-*δ*, *α*-*β*, *α*-*γ*, *β*-*α* and *β*-*γ* CFC. As these decreases involve high-frequency components, we speculate that this finding is likely linked to the characteristic slowing down of signals in AD, i.e. a decrease of high-frequency power [5, 6]. On the other hand, we observe an increase of weighted CBW of *θ*-*θ* WFC and *θ*-*δ* and *γ*-*θ* CFC in AD cases. Interestingly, previously a decrease in *γ*-*θ* phase-amplitude coupling was reported to signify progression from mild cognitive impairment to AD [43]. However, our results indicate an opposite pattern.

Then, weighted and unweighted *V_G_*are used to assess the vulnerability of information integration of the network to the removal of a specific coupling type. Numerical results of comparing unweighted *V_G_* are reported in the Appendix (Table C.XVI) and visualised in Fig. 8. Numerical results of comparing weighted *V_G_* are reported and visualised in Appendix (Table C.XIX) and visualised in Fig. 9.

**FIG. 8:**
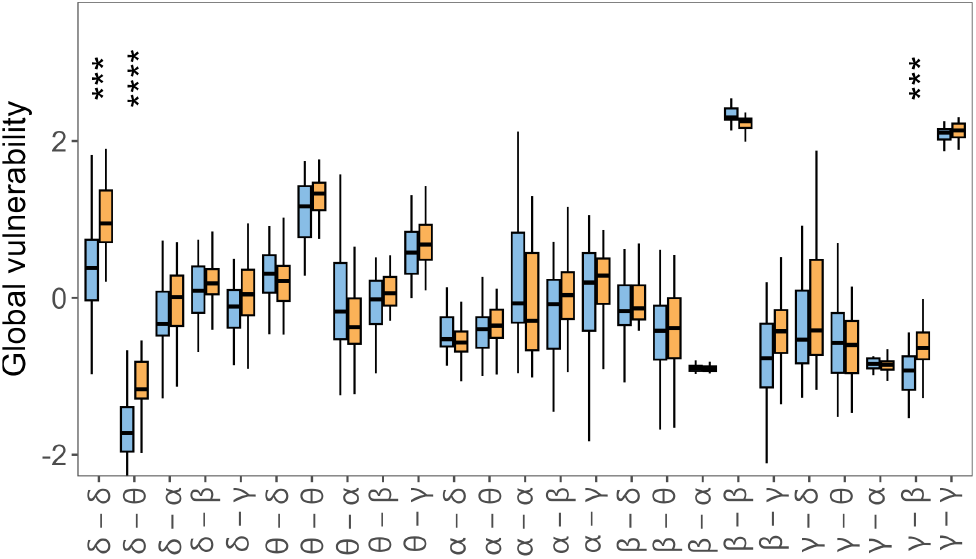
Global vulnerability of HC (blue) and AD (orange). Significant differences observed in at least ten thresholded networks and across all epochs are encoded by asterisks. The number of asterisks corresponds to the p-value (FDR corrected), i.e. *p ≤* 0.0001 “****”, *p ≤* 0.001 “***”, *p ≤* 0.01 “**”, and *p ≤* 0.05 “*”.

**FIG. 9:**
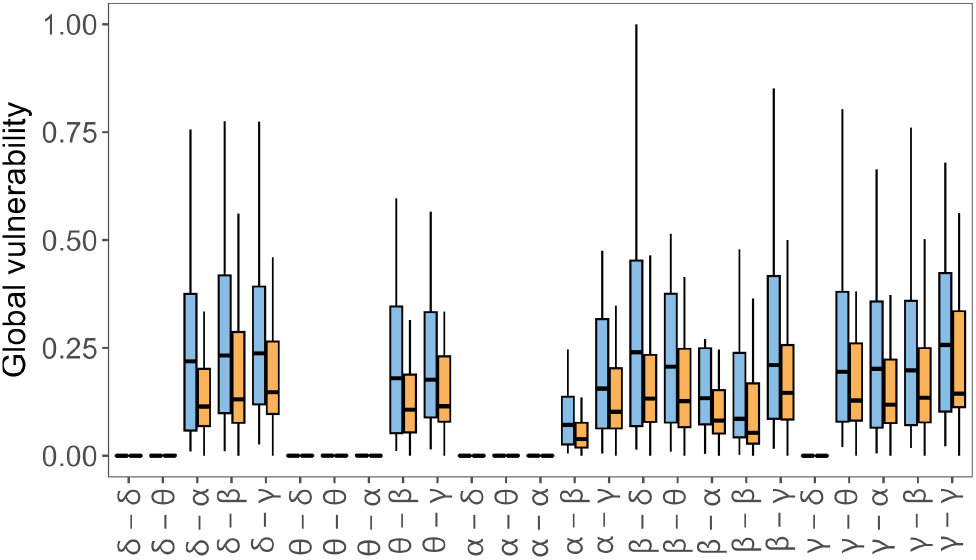
Weighted global vulnerability of HC (blue) and AD (orange). Significant differences observed in at least ten thresholded networks and across all epochs are encoded by asterisks. The number of asterisks corresponds to the p-value (corrected), i.e. *<* 0.001 “****”, 0.001 “***”, 0.01 “**”, and 0.05 “*”.

The AD brain networks seem more vulnerable to removing multiple types of couplings. Weighted *V_G_* fails to detect any reliable differences. We speculate this might be caused by edge weight differences across different coupling types, thus biasing the results. *V_G_*is likely more sensitive towards such an issue, as it is a global measure in contrast to the other measures, which consider predominantly local relationships. A significant increase in unweighted *V_G_* in AD cases is observed in *δ*-*δ* WFC and *δ*-*θ* and *γ*-*β* CFC. Interestingly, the removal of WFC generally causes a larger increase in vulnerability compared to CFC (except for *α*-*α*, suggesting that while CFC plays a crucial role in the brain networks, WFC seems dominant in the brain networks.

Finally, weighted and unweighted *V_L_* are used to assess the vulnerability of segregation of the network to the removal of a particular coupling type. Results of statistical tests comparing unweighted *V_L_* are reported in Appendix (Table C.XVII) and visualised in Fig. 10. Results of statistical tests comparing weighted *V_L_* are reported in Appendix (Table C.XX) and visualised in Fig. 11.

**FIG. 10:**
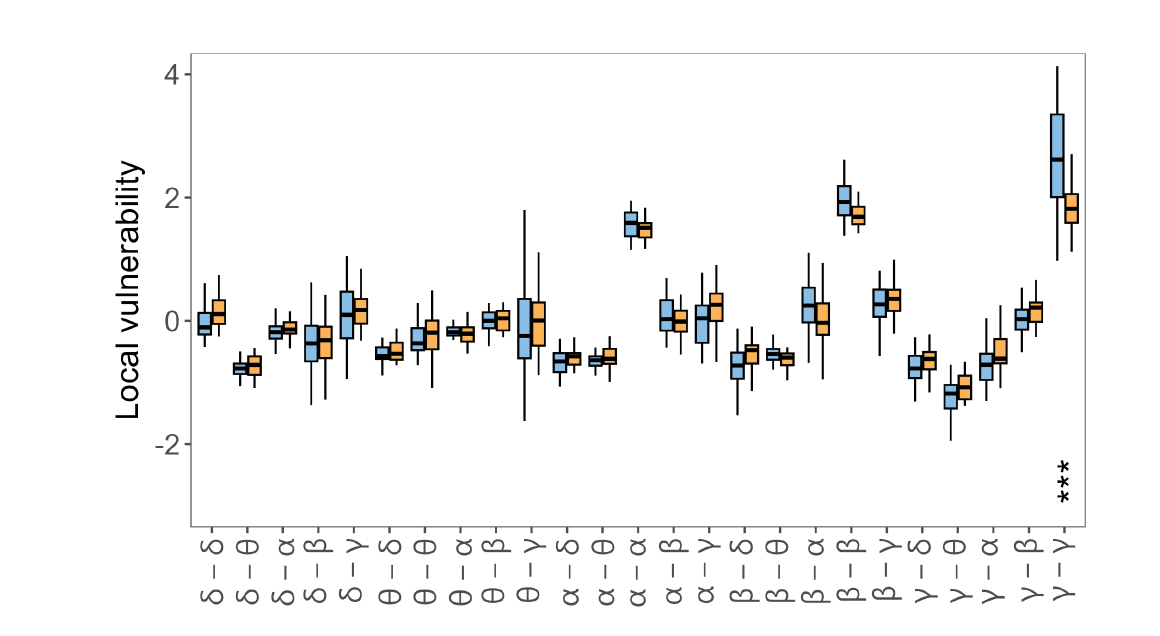
Local vulnerability of HC (blue) and AD (orange). Significant differences observed in at least ten thresholded networks and across all epochs are encoded by asterisks. The number of asterisks corresponds to the p-value (FDR corrected), i.e. *p ≤* 0.0001 “****”, *p ≤* 0.001 “***”, *p ≤* 0.01 “**”, and *p ≤* 0.05 “*”.

**FIG. 11:**
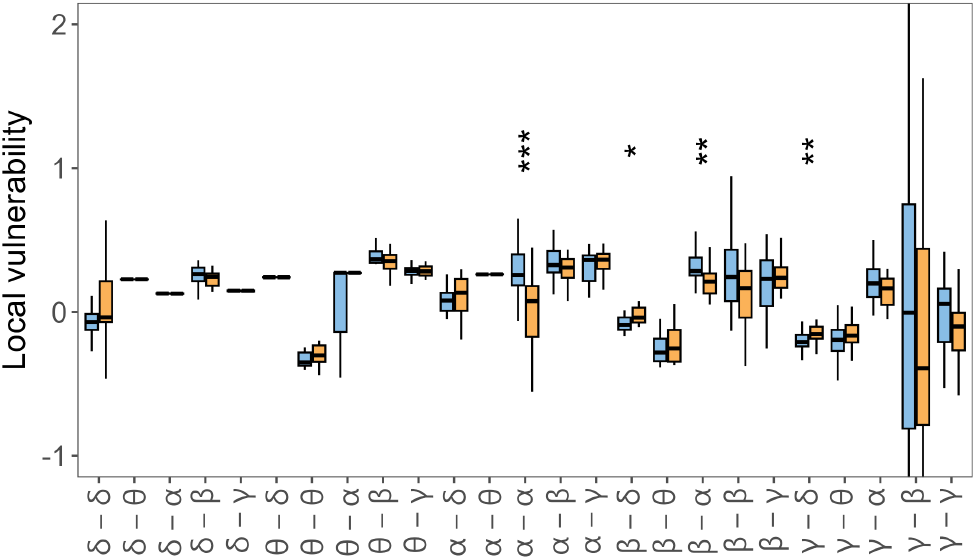
Weighted local vulnerability of HC (blue) and AD (orange). Significant differences observed in at least ten thresholded networks and across all epochs are encoded by asterisks. The number of asterisks corresponds to the p-value (FDR corrected), i.e. *p ≤* 0.0001 “****”, *p ≤* 0.001 “***”, *p ≤* 0.01 “**”, and *p ≤* 0.05 “*”.

*γ*-*γ* is the most robustly linked to segregation measured with unweighted *V_L_*, which fits well with the evidence of high-frequency oscillations being related to local processing [44]. Moreover, this coupling is significantly more vulnerable in weighted networks of HC cases, which is likely related to the decreased *γ* activity in AD [5]. Likely for similar reasons, the removal of *α*-*α* WFC and *β*-*α* CFC causes a significant increase of weighted *V_L_* in HC, suggesting the segregation function enabled by these high-frequency components is likely disrupted in AD. On the other hand, *β*-*δ* and *γ*-*δ* CFC removal cause a significant increase of weighted *V_L_*in AD cases. This suggests that in AD cases, the high-frequency CFC takes over the role of enabling network segregation as the high-frequency WFC is attenuated.

### D. Classification results

SVM classifiers were trained using network features extracted from CS and CBS separately to measure the predictive power of CS and CBS-based networks and evaluate the multilayer network features (Fig. 12). A detailed performance summary of the best models is reported in Table I.

**FIG. 12:**
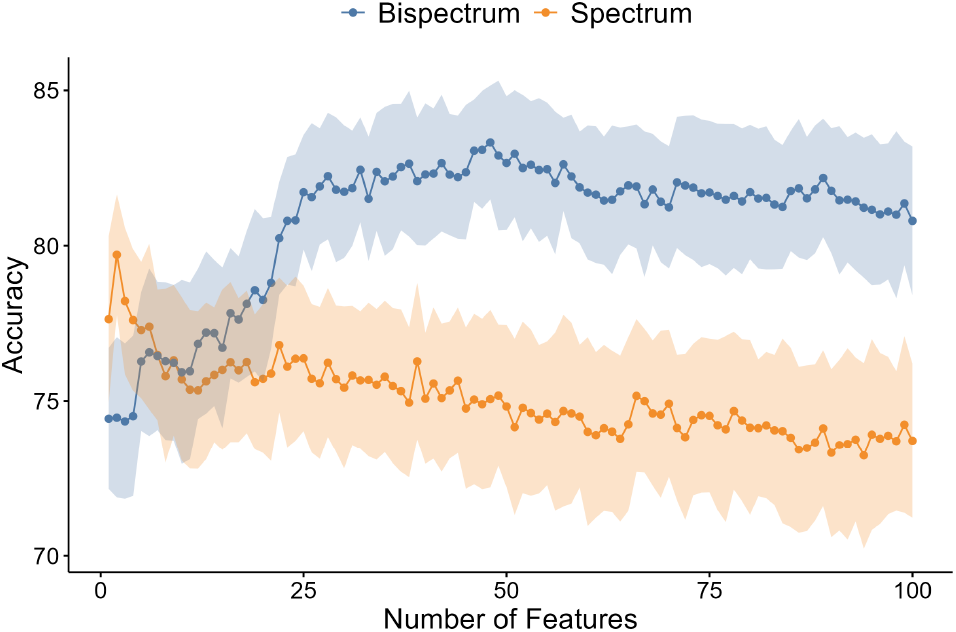
Average accuracy (points) with standard deviation (ribbons) of the classifiers trained with graph-theory features using a 10-fold stratified cross-validation repeated 100 times. Specifically, the features considered are node strength for cross-spectrum (orange) and node strength, CBW, *V_G_* and *V_L_* cross-bispectrum (blue) networks. The features are sequentially added to the classifier based on their effect size.

**TABLE I:** Performance of the best models trained using network features identified via forward feature selection. The feature sets contain 2, 48 and 50 features for the spectrum, bispectrum and combined models, respectively.

| Model | Accuracy | Sensitivity | Specificity |
| --- | --- | --- | --- |
| Spectrum | 79.71 % (SD=1.94) | 86.42 % (SD=2.95) | 74.62 % (SD=4.48) |
| Bispectrum | 83.32 % (SD=1.83) | 86.62 % (SD=2.95) | 80.71 % (SD=3.9) |
| Combined | 81.39 % (SD=2.09) | 85.97 % (SD=2.45) | 78.91 % (SD=3.28) |

All CBS-based models outperform the CS-based models, suggesting that information related to nonlinear and CFC coupling might be crucial for the modelling and classification of AD. However, such a conclusion might be biased as the CS-based models are trained using a smaller set of features, i.e. node strengths. Thus, the comparison is likely unfair and should be interpreted conservatively. The best CS-based model reaches its highest accuracy of 79.71% (SD=1.94) using only two features. These features are the node strengths of channels F4-C4 and C3-P3 in the *θ* frequency band WFC. In contrast, the CBS-based models require considerably more features to achieve the highest accuracy of 83.32% (SD=1.83) with 48 features (Table II). Interestingly, the majority of these features are CFC. Furthermore, the weighted CBW seems to provide the most information to the classifier from the multilayer network measures introduced in this study, as it is included multiple times in the final feature set. Node strengths from all areas are utilised, but the central-parietal channels are selected repeatedly across multiple frequency couplings. It is worth noting that including the same features across different network thresholds appears to improve the performance, despite likely strong correlations between such features. If only a single network threshold was selected (i.e. based on the largest effect size), the accuracy drops by 2%-3%. Interestingly, both CS- and CBS-based models utilise the F4-C4 channel from the *θ* WFC, suggesting some shared information between these two functional connectivity methods.

**TABLE II:** Features included in the best cross-bispectrum-based classifier. For multilayer network metrics, the network thresholds are in parentheses. This is not necessary for the node strengths as these are obtained from the unthresholded networks.

| coupling | Node strength | Multilayer network metric |
| --- | --- | --- |
| α-β | C4P4, CZPZ, P3O1, T5O1 | |
| β-α | C4P4, P3O1 | weighted CBW (0.7) |
| δ-α | F4C4 | weighted CBW (0.4, 0.45, 0.5, 0.55, 0.6) |
| δ-θ | C3P3, C4P4, CZPZ, FZCZ, O1O2 | $V_G$ (0.55) |
| γ-α | F4FZ | weighted CBW (0.4, 0.45, 0.5, 0.55, 0.6, 0.65, 0.7, 0.75, 0.8, 0.85) |
| θ-δ | CZPZ, FZCZ, P4PZ | |
| θ-θ | F4C4 | |
| δ-δ | | $V_G$ (0.65) |
| α-δ | | weighted CBW (0, 0.05, 0.1, 0.15, 0.2, 0.25) |
| α-γ | | weighted CBW (0.5, 0.55, 0.6, 0.65, 0.7, 0.75, 0.8) |

Finally, we trained a combined model with the sets of best features concatenated from the CS- and CBS-based models, i.e. using 50 features. However, the accuracy of such a model is only 81.39% (SD=2.09), which is lower than the CBS-based model suggesting that the addition of CS-based features introduces redundant information into the model.

## V. CONCLUSIONS AND FUTURE WORK

We have demonstrated that CBS and CS detect similar differences between AD and HC networks, but CBS has an advantage over CS by including cross-frequency and nonlinear interactions. We report several significant differences in CFC both globally and on a node level, suggesting that including CFC in a graph-theoretic analysis of brain networks is crucial to obtain a more detailed insight into their structure and function. Furthermore, we show that multilayer network analysis provides a simple yet powerful framework for representing and analysing the role of CFC in brain networks. Using this framework, we present a novel approach to elucidate the roles of different frequency components of EEG signals. Moreover, we show that both CFC and WFC CBS-based networks can be used to classify AD with high accuracy.

CFC has been suggested to be related to modulatory activity, i.e. slow band modulating the activity of fast oscillations. However, it remains unclear why low-frequency CFC would be increased in AD and requires further in-depth study.

Next, although (cross-)bispectrum was shown to be a powerful tool to detect various types of WFC and CFC, such as phase-phase or phase-amplitude, CBS seems to capture an unknown mixture of these types of couplings. Therefore, a combination of bispectrum with other types of CFC methods might be a plausible direction for future research.

Furthermore, by relying on traditional frequency bands to define the layers of the networks, our framework might miss some CFC occurring on finer scales, e.g. interaction within one band. However, considering the CFC within only a few bands allows us to construct multilayer networks with a relatively small number of layers. Thus, we argue that relying on the five bands is necessary to introduce the CFC into network analysis without increasing the complexity significantly.

The presented multilayer network analysis focused only on how dependent or vulnerable the networks are on different types of frequency coupling to enable integration and segregation properties. Although these two properties are hypothesised to be crucial in brain networks, their analysis might be insufficient to elucidate the function of the frequency couplings. Thus, we suggest to focus on other graph-theoretic measures beyond integration and segregation in future work. Although these two properties are hypothesised to be crucial in brain networks, their analysis is not sufficient to elucidate the functions the frequency couplings might enable across various spatio-temporal scales in normal brains and how these functions disappear or change in AD.

A limitation of our study is the relatively small sample size. This leads to some of the observed significant differences being underpowered. Thus, the small differences we report in this study should be interpreted more conservatively. However, despite this limitation, we identify a set of reliable biomarkers as evidenced by the classification results. In future research, it might also be important and interesting to explore more complex graph-based features that would capture the differences between AD and HC in a lower-dimensional space more efficiently.

## ACKNOWLEDGMENT

The EEG data was funded by a grant from the Alzheimer’s Research UK (ARUK-PPG20114B-25). The views expressed are those of the author(s) and not necessarily those of the NHS, the NIHR or the Department of Health.

## Appendix A: Statistical power

We calculate the statistical power given our sample size (*N* = 40) as a function of the effect size (Cohen’s d) of a two-sample t-test with a two-sided alternative hypothesis and a significant p-value of 0.05 (Figure A.1). Sufficient power (80%) is reached with an effect size ≥ 0.45.

**FIG. A.1:**
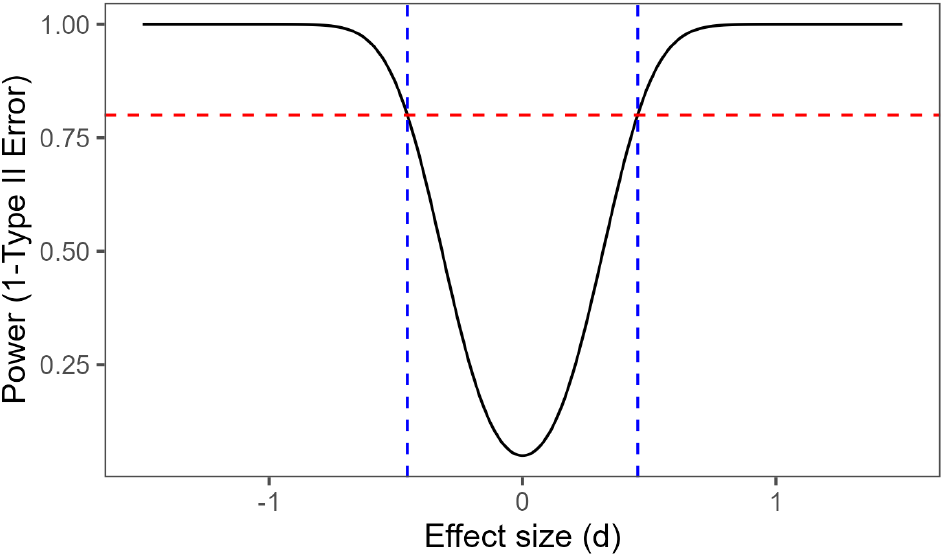
Statistical power for different values of Cohen’s d effect size with sample size 40.

## Appendix B: Results for epoch 1

In this section, we report the results of the statistical comparisons and corresponding figures using only data from the first epoch.

The comparisons of average coupling computed with cross-spectrum and cross-bispectrum are shown in Tables B.I and B.II, respectively. Additionally, the corresponding figures of connectivity matrices computed with cross-spectrum and cross-bispectrum are shown in Figures B.2 and B.3.

The comparisons of node strength computed with cross-spectrum and cross-bispectrum are shown in Table B.III and Figure B.4, and Table B.IV and Figure B.5, respectively.

Results of comparisons of the unweighted multilayer network metrics are reported in Tables B.V, B.VI and B.VII for edge betweenness, global vulnerability and local vulnerability, respectively. The corresponding figures are; edge betweenness (Figure B.6), global vulnerability (Figure B.7) and local vulnerability (Figure B.8).

Results of comparisons of the weighted multilayer network metrics are reported in Tables B.VIII, B.IX and B.X for edge betweenness, global vulnerability and local vulnerability, respectively. The corresponding figures are; edge betweenness (Figure B.9), global vulnerability (Figure B.10) and local vulnerability (Figure B.11).

**FIG. B.2:**
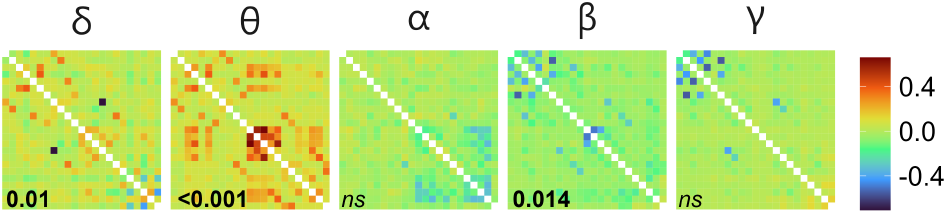
Difference of average connectivity matrices (*AD − HC*) measured with cross-spectrum in epoch 1 of (A) AD and (B) HC. For visualisation purposes, the values were min-max normalised. Digits in black denote a *p*-value testing for the difference in global coupling (*p <* 0.05 in bold, in italics otherwise).

**TABLE B.I:**
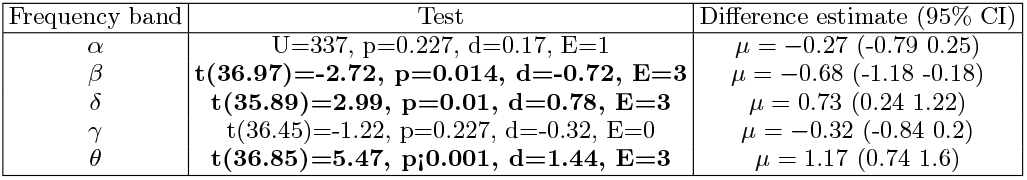
Comparisons of the mean of adjacency matrix constructed with cross-spectrum in epoch 1. The results are reported as follows: statistics value (degrees of freedom), p-value of the test, Cohen’s d effect size (or nonparametric alternative), number of epochs where significant differences were observed (E), difference estimate *µ* with 95% confidence interval. Reliable differences (significant in all three epochs) are highlighted with bold text.

| Frequency band | Test | Difference estimate (95% CI) |
| --- | --- | --- |
| $\alpha$ | $U=337, p=0.227, d=0.17, E=1$ | $\mu = -0.27 (-0.79 \ 0.25)$ |
| $\beta$ | **$t(36.97)=-2.72, p=0.014, d=-0.72, E=3$** | $\mu = -0.68 (-1.18 \ -0.18)$ |
| $\delta$ | **$t(35.89)=2.99, p=0.01, d=0.78, E=3$** | $\mu = 0.73 (0.24 \ 1.22)$ |
| $\gamma$ | $t(36.45)=-1.22, p=0.227, d=-0.32, E=0$ | $\mu = -0.32 (-0.84 \ 0.2)$ |
| $\theta$ | **$t(36.85)=5.47, p<0.001, d=1.44, E=3$** | $\mu = 1.17 (0.74 \ 1.6)$ |

**FIG. B.3:**
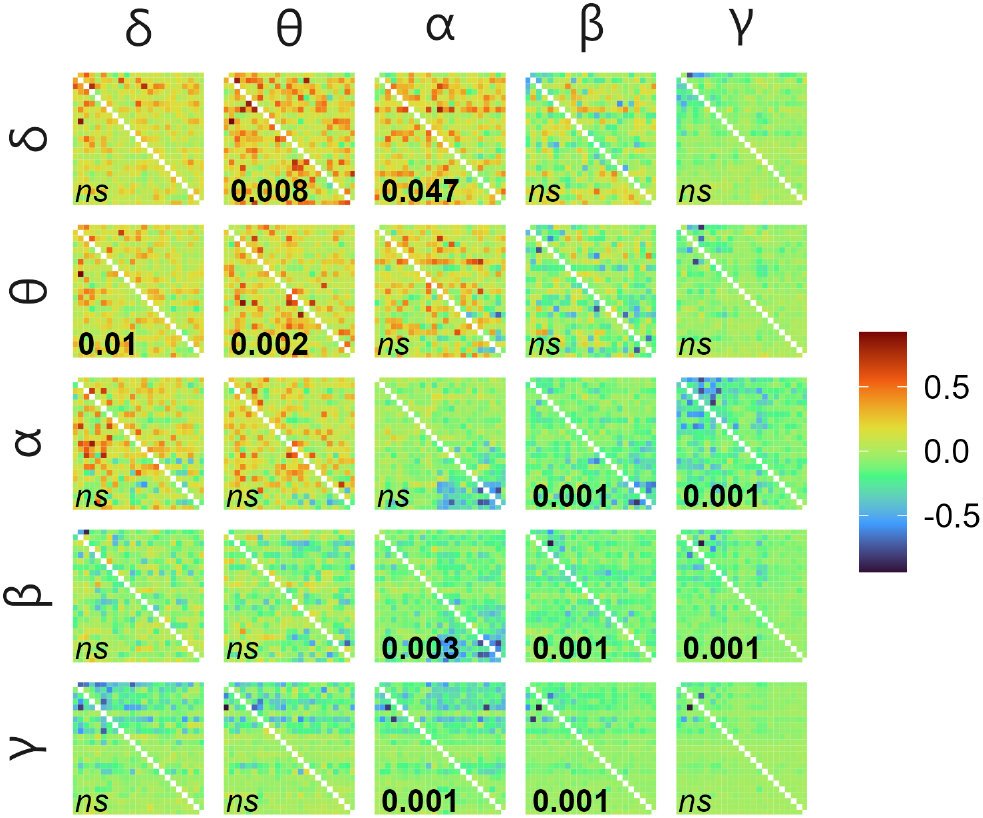
Difference of average connectivity matrices (*AD − HC*) measured with cross-bispectrum with input frequency on the vertical facets and output frequency on the horizontal. For visualisation purposes, the values were min-max normalised. Digits in black denote a *p*-value testing for the difference in global coupling (*p <* 0.05 in bold, in italics otherwise).

**TABLE B.II:**
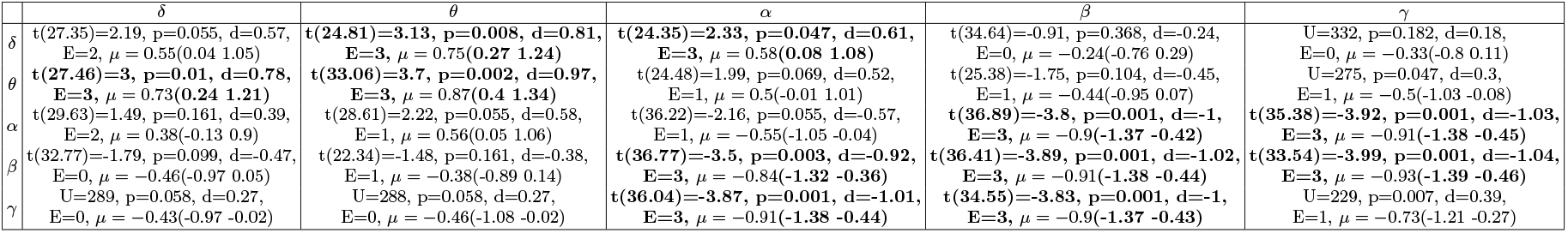
Comparisons of the mean of adjacency matrix constructed with cross-bispectrum in epoch 1. The results are reported as follows: statistics value (degrees of freedom), p-value of the test, Cohen’s d effect size (or nonparametric alternative), number of epochs where significant differences were observed (E), difference estimate *µ* with 95% confidence interval. Reliable differences (significant in all three epochs) are highlighted with bold text.

**FIG. B.4:**
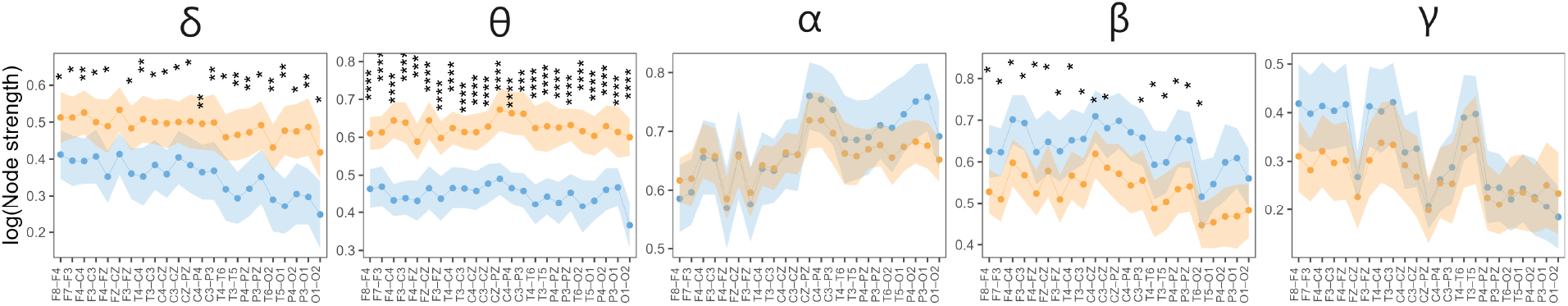
Node strength (min-max normalised) measured with CS in epoch 1 of HC (blue) and AD (orange): mean with 95% confidence intervals. Significant differences (*p ≤* 0.05) observed in at least ten thresholded networks are encoded by asterisks. The number of asterisks corresponds to the p-value (FDR corrected), i.e. *p ≤* 0.0001 “****”, *p ≤* 0.001 “***”, *p ≤* 0.01 “**”, and *p ≤* 0.05 “*”.

**TABLE B.III:**
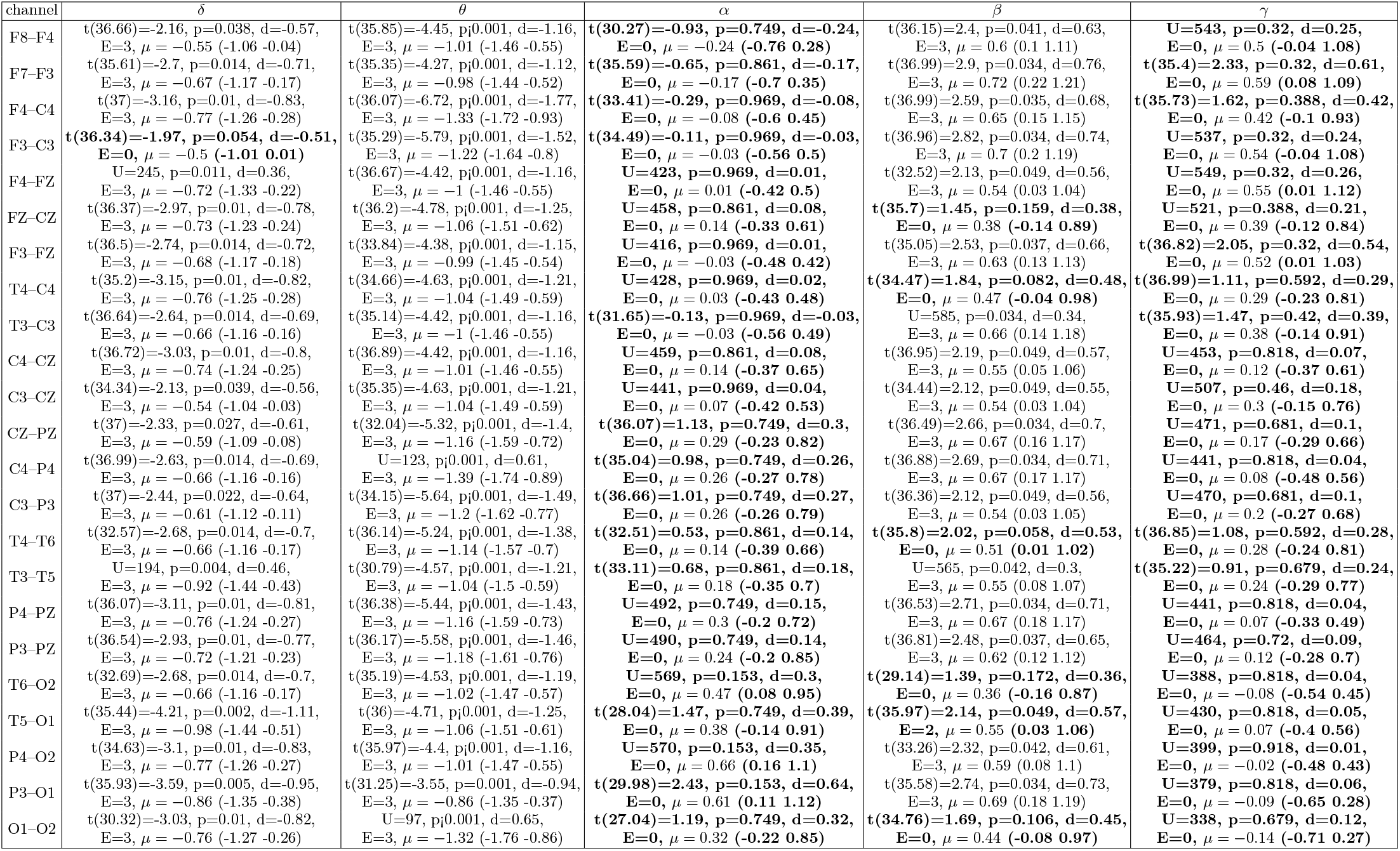
Comparisons of node strength measured with cross-spectrum in epoch 1. The results are reported as follows: statistics value (degrees of freedom), p-value of the test, Cohen’s d effect size (or nonparametric alternative), number of epochs where significant differences were observed (E), and difference estimate *µ* with 95% confidence interval (CI). Reliable differences (significant in all three epochs) are highlighted with bold text.

**FIG. B.5:**
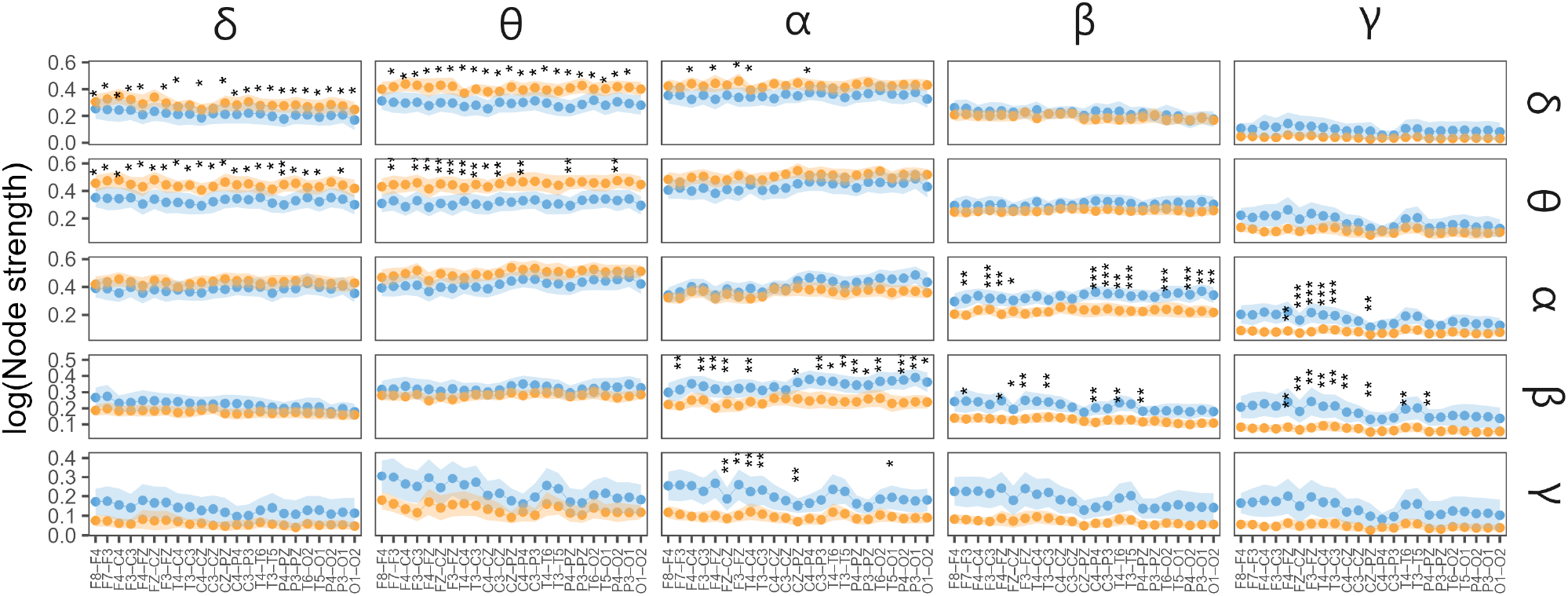
Node strength (min-max normalised) measured with CBS in epoch 1 of HC (blue) and AD (orange): mean with 95% confidence intervals. The input frequency is on the vertical facets, and the output frequency is on the horizontal. Significant differences (*p ≤* 0.05) observed in at least ten thresholded networks are encoded by asterisks. The number of asterisks corresponds to the p-value (FDR corrected), i.e. *p ≤* 0.0001 “****”, *p ≤* 0.001 “***”, *p ≤* 0.01 “**”, and *p ≤* 0.05 “*”.

**TABLE B.IV:**
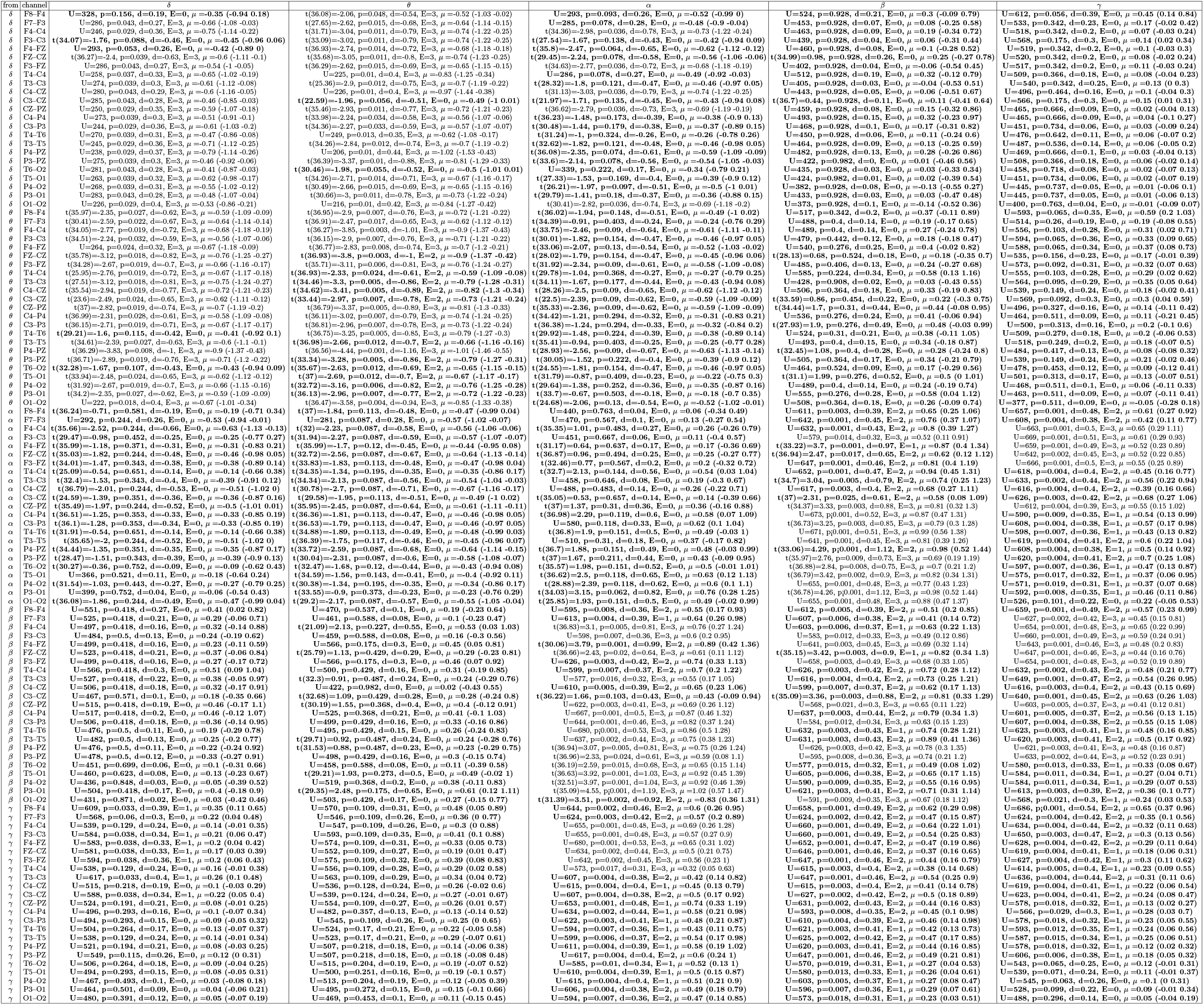
Comparisons of node strength measured with cross-bispectrum in epoch 1. The results are reported as follows: statistics value (degrees of freedom), p-value of the test, Cohen’s d effect size (or nonparametric alternative), number of epochs where significant differences were observed (E), and difference estimate *µ* with 95% confidence interval (CI). Reliable differences (significant in all three epochs) are highlighted with bold text.

**TABLE B.V:** Results from epoch 1 comparing unweighted edge betweenness. The results are reported as follows: statistics value (degrees of freedom), p-value of the test, Cohen’s d effect size or nonparameteric alternative, number of thresholds where significant differences were observed (T), number of epochs where significant differences were observed (E), group difference estimate (95% CI). Reliable differences (significant in all three epochs) are highlighted with bold text.

| | $\delta$ | $\theta$ | $\alpha$ | $\beta$ | $\gamma$ |
| --- | --- | --- | --- | --- | --- |
| $\delta$ | t(36.68)=0.95, p=0.348, d=0.25,<br>T=1, E=0, $\mu = 0.25$ (-0.28 0.77) | U=508, p=0.175, d=0.18,<br>T=0, E=0, $\mu = 0.31$ (-0.15 0.74) | U=591, p=0.007, d=0.35,<br>T=16, E=2, $\mu = 0.62$ (0.19 1.09) | **U=636, p=0.001, d=0.44,**<br>**T=17, E=3, $\mu = 0.7$ (0.33 1.1)** | U=542, p=0.058, d=0.25,<br>T=9, E=0, $\mu = 0.32$ (0 0.54) |
| $\theta$ | U=455, p=0.594, d=0.07,<br>T=0, E=0, $\mu = 0.11$ (-0.28 0.55) | U=513, p=0.151, d=0.19,<br>T=3, E=0, $\mu = 0.28$ (-0.15 0.71) | U=500, p=0.218, d=0.16,<br>T=0, E=0, $\mu = 0.28$ (-0.15 0.76) | U=517, p=0.134, d=0.2,<br>T=0, E=0, $\mu = 0.33$ (-0.11 0.73) | U=498, p=0.23, d=0.16,<br>T=0, E=0, $\mu = 0.2$ (-0.13 0.57) |
| $\alpha$ | t(36.86)=1.34, p=0.185, d=0.35,<br>T=0, E=0, $\mu = 0.35$ (-0.17 0.87) | t(30.26)=1.16, p=0.25, d=0.3,<br>T=0, E=0, $\mu = 0.3$ (-0.22 0.82) | U=483, p=0.333, d=0.13,<br>T=0, E=0, $\mu = 0.18$ (-0.3 0.64) | U=521, p=0.118, d=0.21,<br>T=0, E=0, $\mu = 0.31$ (-0.1 0.73) | U=528, p=0.095, d=0.22,<br>T=3, E=0, $\mu = 0.19$ (-0.04 0.42) |
| $\beta$ | U=489, p=0.289, d=0.14,<br>T=0, E=0, $\mu = 0.19$ (-0.15 0.54) | U=475, p=0.399, d=0.11,<br>T=0, E=0, $\mu = 0.15$ (-0.23 0.59) | U=472, p=0.426, d=0.11,<br>T=0, E=0, $\mu = 0.18$ (-0.25 0.66) | U=494, p=0.255, d=0.15,<br>T=1, E=0, $\mu = 0.23$ (-0.18 0.66) | U=508, p=0.175, d=0.18,<br>T=1, E=0, $\mu = 0.27$ (-0.14 0.6) |
| $\gamma$ | U=475, p=0.399, d=0.11,<br>T=0, E=0, $\mu = 0.15$ (-0.22 0.52) | U=504, p=0.195, d=0.17,<br>T=0, E=0, $\mu = 0.2$ (-0.13 0.53) | U=548, p=0.047, d=0.26,<br>T=11, E=2, $\mu = 0.25$ (0.01 0.48) | U=479, p=0.365, d=0.12,<br>T=0, E=0, $\mu = 0.16$ (-0.18 0.44) | U=360, p=0.357, d=0.12,<br>T=0, E=0, $\mu = -0.22$ (-0.68 0.23) |

**TABLE B.VI:**
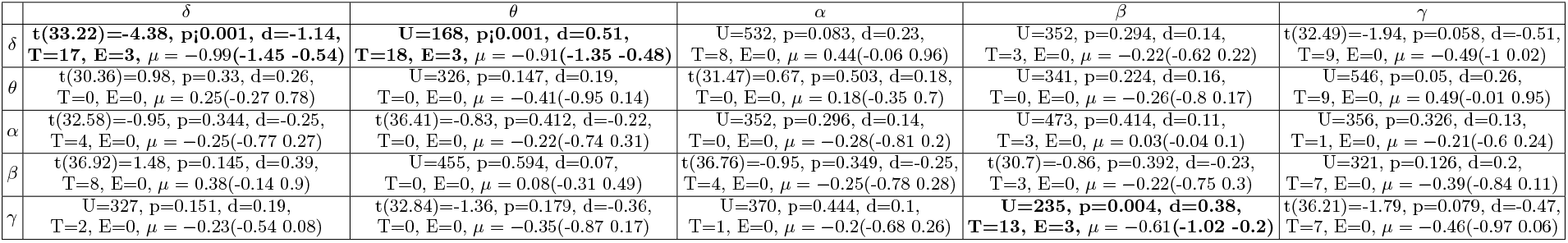
Results from epoch 1 comparing unweighted global vulnerability. The results are reported as follows: statistics value (degrees of freedom), p-value of the test, Cohen’s d effect size or nonparameteric alternative, number of thresholds where significant differences were observed (T), number of epochs where significant differences were observed (E), group difference estimate (95% CI). Reliable differences (significant in all three epochs) are highlighted with bold text.

**TABLE B.VII:** Results from epoch 1 comparing unweighted local vulnerability. The results are reported as follows: statistics value (degrees of freedom), p-value of the test, Cohen’s d effect size or nonparameteric alternative, number of thresholds where significant differences were observed (T), number of epochs where significant differences were observed (E), group difference estimate (95% CI). Reliable differences (significant in all three epochs) are highlighted with bold text.

| | $\delta$ | $\theta$ | $\alpha$ | $\beta$ | $\gamma$ |
| --- | --- | --- | --- | --- | --- |
| $\delta$ | U=308, p=0.083, d=0.23,<br>T=9, E=0, $\mu = -0.4$ (-0.82 0.06) | t(36.15)=-0.87, p=0.389, d=-0.23,<br>T=2, E=0, $\mu = -0.23$ (-0.75 0.3) | t(37)=1.86, p=0.068, d=0.49,<br>T=9, E=0, $\mu = 0.48$ (-0.04 0.99) | t(35.43)=-1.95, p=0.056, d=-0.51,<br>T=7, E=0, $\mu = -0.5$ (-1.01 0.01) | t(34.87)=-1.11, p=0.27, d=-0.29,<br>T=0, E=0, $\mu = -0.29$ (-0.81 0.23) |
| $\theta$ | U=361, p=0.365, d=0.12,<br>T=0, E=0, $\mu = -0.21$ (-0.65 0.26) | t(32.16)=-0.52, p=0.606, d=-0.14,<br>T=2, E=0, $\mu = -0.14$ (-0.66 0.39) | U=389, p=0.638, d=0.06,<br>T=0, E=0, $\mu = -0.08$ (-0.45 0.29) | t(29.01)=-0.85, p=0.399, d=-0.22,<br>T=0, E=0, $\mu = -0.22$ (-0.74 0.3) | t(36.97)=-0.56, p=0.581, d=-0.15,<br>T=0, E=0, $\mu = -0.15$ (-0.67 0.38) |
| $\alpha$ | t(35.69)=-1.77, p=0.082, d=-0.46,<br>T=9, E=0, $\mu = -0.45$ (-0.97 0.06) | t(36.82)=-0.42, p=0.677, d=-0.11,<br>T=0, E=0, $\mu = -0.11$ (-0.64 0.42) | U=504, p=0.195, d=0.17,<br>T=5, E=0, $\mu = 0.3$ (-0.19 0.61) | t(29.31)=1.25, p=0.216, d=0.33,<br>T=3, E=0, $\mu = 0.32$ (-0.2 0.84) | t(34.1)=-1.23, p=0.223, d=-0.32,<br>T=0, E=0, $\mu = -0.32$ (-0.84 0.2) |
| $\beta$ | t(36.26)=-1.61, p=0.114, d=-0.42,<br>T=5, E=0, $\mu = -0.42$ (-0.94 0.1) | U=360, p=0.357, d=0.12,<br>T=0, E=0, $\mu = -0.27$ (-0.74 0.25) | t(36.89)=-0.95, p=0.344, d=-0.25,<br>T=2, E=0, $\mu = -0.25$ (-0.77 0.27) | U=519, p=0.126, d=0.2,<br>T=1, E=0, $\mu = 0.39$ (-0.08 0.96) | t(23.65)=0.83, p=0.409, d=0.22,<br>T=4, E=0, $\mu = 0.22$ (-0.31 0.74) |
| $\gamma$ | U=283, p=0.033, d=0.28,<br>T=11, E=1, $\mu = -0.4$ (-0.81 -0.03) | t(32.95)=-2.29, p=0.026, d=-0.6,<br>T=11, E=1, $\mu = -0.58$ (-1.08 -0.07) | t(36.03)=-1.93, p=0.059, d=-0.51,<br>T=9, E=0, $\mu = -0.5$ (-1.01 0.02) | U=474, p=0.408, d=0.11,<br>T=2, E=0, $\mu = 0.16$ (-0.26 0.61) | **U=635, p=0.001, d=0.44,**<br>**T=13, E=3, $\mu = 0.3$ (0.12 0.49)** |

**TABLE B.VIII:**
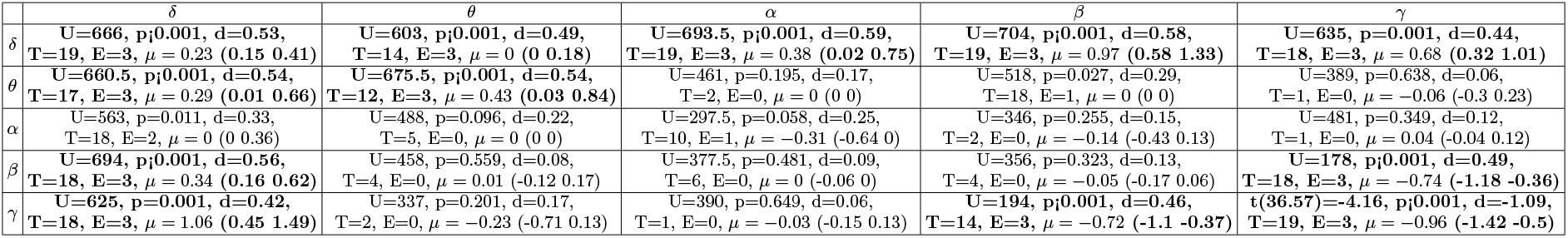
Results from epoch 1 comparing weighted edge betweenness. The results are reported as follows: statistics value (degrees of freedom), p-value of the test, Cohen’s d effect size or nonparameteric alternative, number of thresholds where significant differences were observed (T), number of epochs where significant differences were observed (E), group difference estimate (95% CI). Reliable differences (significant in all three epochs) are highlighted with bold text.

**TABLE B.IX:**
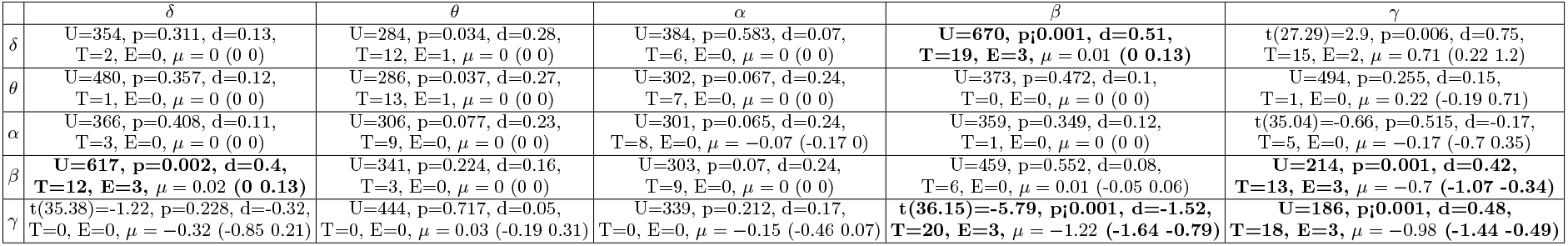
Results from epoch 1 comparing weighted global vulnerability. The results are reported as follows: statistics value (degrees of freedom), p-value of the test, Cohen’s d effect size or nonparameteric alternative, number of thresholds where significant differences were observed (T), number of epochs where significant differences were observed (E), group difference estimate (95% CI). Reliable differences (significant in all three epochs) are highlighted with bold text.

**TABLE B.X:**
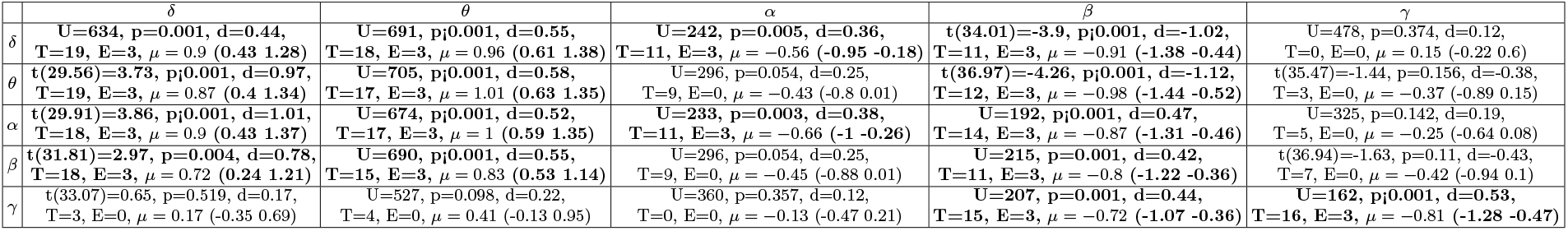
Results from epoch 1 comparing weighted local vulnerability. The results are reported as follows: statistics value (degrees of freedom), p-value of the test, Cohen’s d effect size or nonparameteric alternative, number of thresholds where significant differences were observed (T), number of epochs where significant differences were observed (E), group difference estimate (95% CI). Reliable differences (significant in all three epochs) are highlighted with bold text.

**FIG. B.6:**
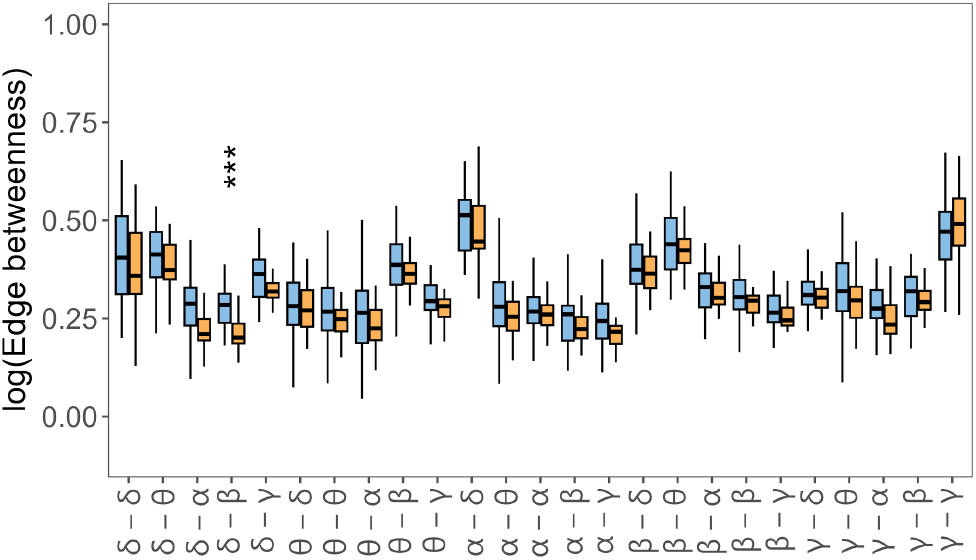
Importance of each type of frequency coupling of HC (blue) and AD (orange) measured by edge betweenness in epoch 1. Significant differences (*p ≤* 0.05) observed in at least ten thresholded networks are encoded by asterisks. The number of asterisks corresponds to the p-value (FDR corrected), i.e. *p ≤* 0.0001 “****”, *p ≤* 0.001 “***”, *p ≤* 0.01 “**”, and *p ≤* 0.05 “*”.

**FIG. B.7:**
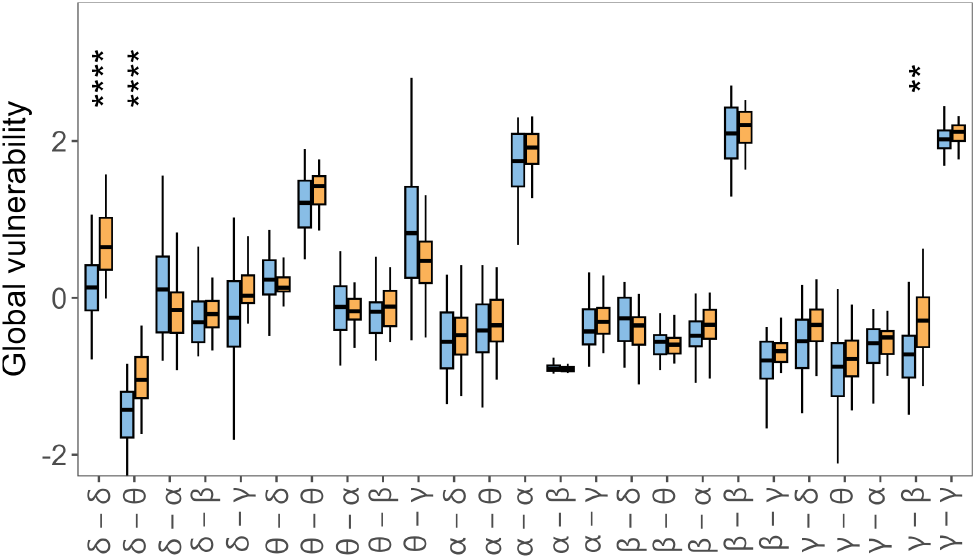
Global vulnerability of HC (blue) and AD (orange) in epoch 1. Significant differences (*p ≤* 0.05) observed in at least ten thresholded networks are encoded by asterisks. The number of asterisks corresponds to the p-value (FDR corrected), i.e. *p ≤* 0.0001 “****”, *p ≤* 0.001 “***”, *p ≤* 0.01 “**”, and *p ≤* 0.05 “*”.

**FIG. B.8:**
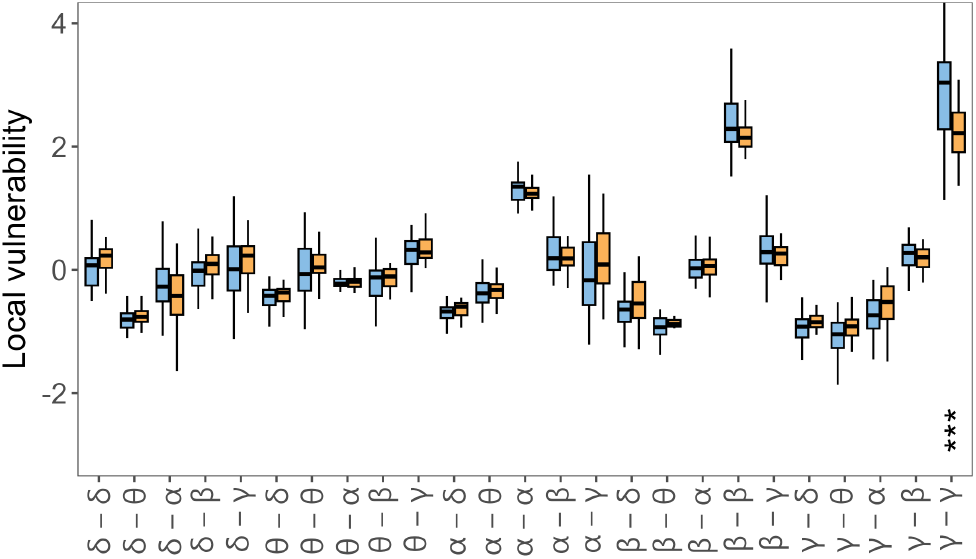
Local vulnerability of HC (blue) and AD (orange) in epoch 1. Significant differences (*p ≤* 0.05) observed in at least ten thresholded networks are encoded by asterisks. The number of asterisks corresponds to the p-value (FDR corrected), i.e. *p ≤* 0.0001 “****”, *p ≤* 0.001 “***”, *p ≤* 0.01 “**”, and *p ≤* 0.05 “*”.

**FIG. B.9:**
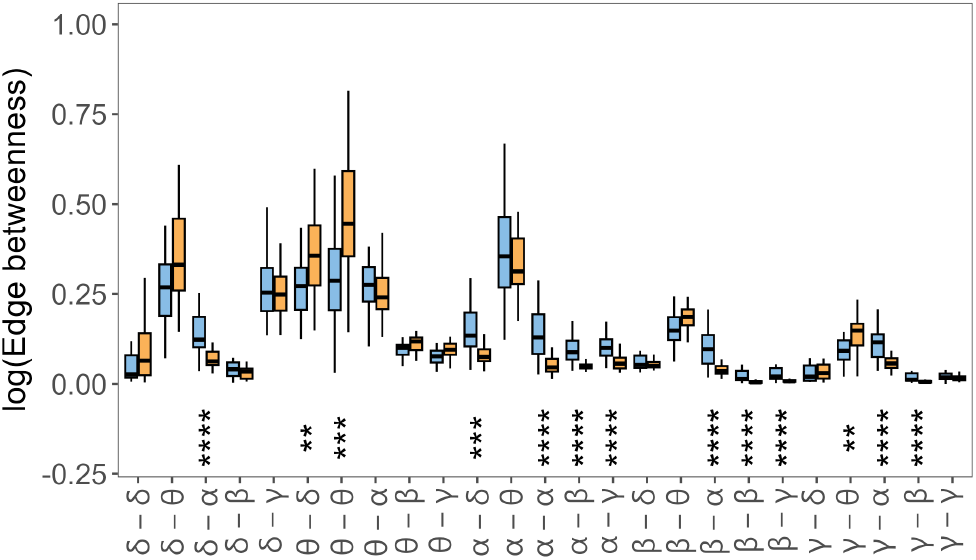
Importance of each type of frequency coupling of HC (blue) and AD (orange) measured by weighted edge betweenness in epoch 1. Significant differences (*p ≤* 0.05) observed in at least ten thresholded networks are encoded by asterisks. The number of asterisks corresponds to the p-value (FDR corrected), i.e. *p ≤* 0.0001 “****”, *p ≤* 0.001 “***”, *p ≤* 0.01 “**”, and *p ≤* 0.05 “*”.

**FIG. B.10:**
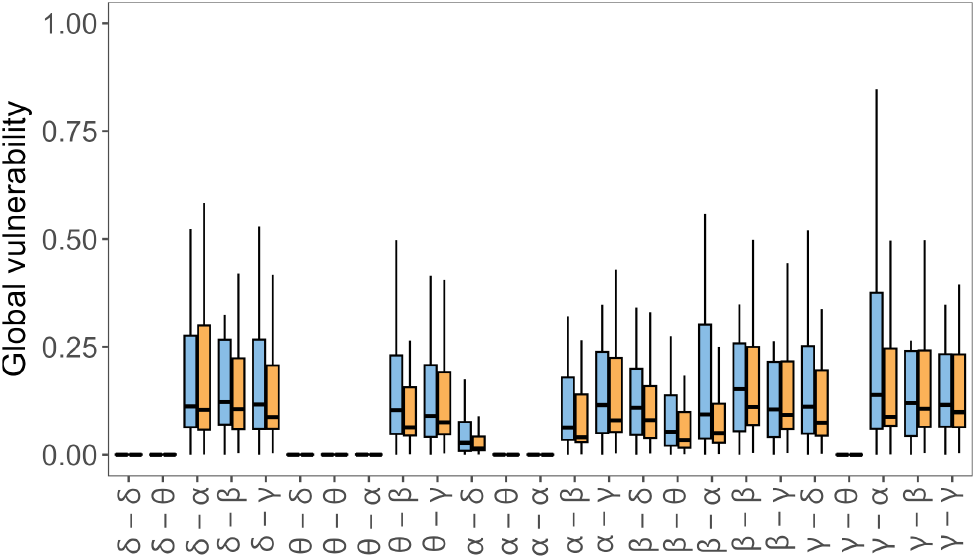
Weighted global vulnerability of HC (blue) and AD (orange) in epoch 1. Significant differences (*p ≤* 0.05) observed in at least ten thresholded networks are encoded by asterisks. The number of asterisks corresponds to the p-value (FDR corrected), i.e. *p ≤* 0.0001 “****”, *p ≤* 0.001 “***”, *p ≤* 0.01 “**”, and *p ≤* 0.05 “*”.

**FIG. B.11:**
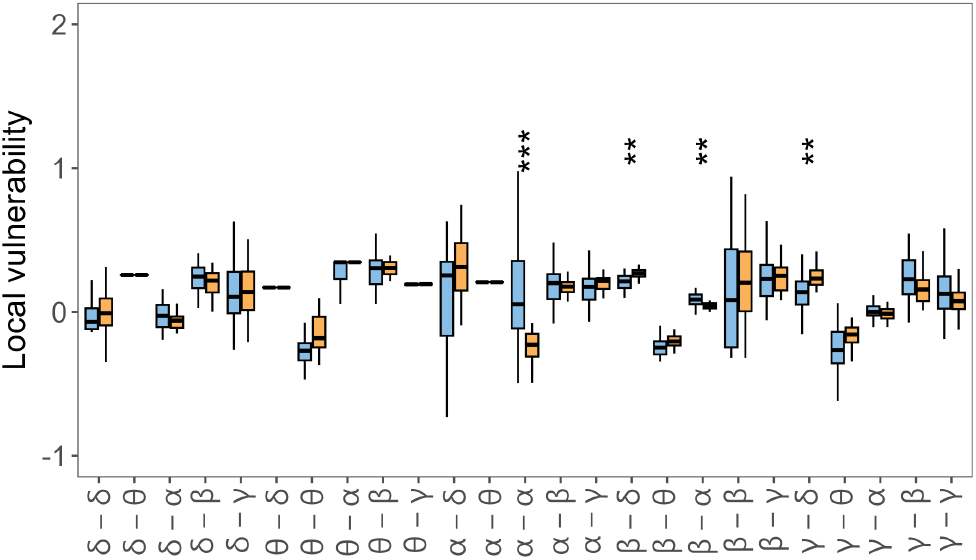
Weighted local vulnerability of HC (blue) and AD (orange) in epoch 1. Significant differences (*p ≤* 0.05) observed in at least ten thresholded networks are encoded by asterisks. The number of asterisks corresponds to the p-value (FDR corrected), i.e. *p ≤* 0.0001 “****”, *p ≤* 0.001 “***”, *p ≤* 0.01 “**”, and *p ≤* 0.05 “*”.

## Appendix C: Results of statisitcal tests for epoch 2

In this section, we report the detailed results of the statistical comparisons accompanying the figures reported in the main text computed using only data from the second epoch. The comparisons of average coupling computed with cross-spectrum and cross-bispectrum are shown in Tables C.XI and C.XII, respectively. The comparisons of node strength computed with cross-spectrum and cross-bispectrum are shown in Tables C.XIII and C.XIV, respectively. Results of comparisons of the unweighted multilayer network metrics are reported in Tables C.XV, C.XVI and C.XVII for edge betweenness, global vulnerability and local vulnerability, respectively. Results of comparisons of the weighted multilayer network metrics are reported in Tables C.XVIII, C.XIX and C.XX for edge betweenness, global vulnerability and local vulnerability, respectively.

**TABLE C.XI:**
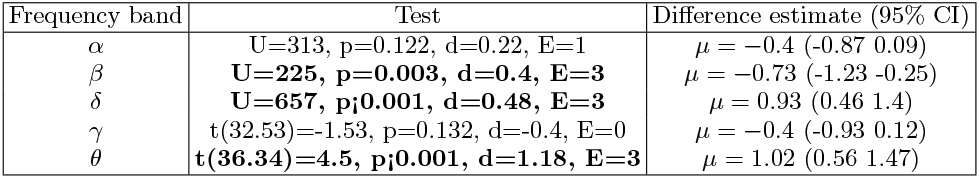
Comparisons of the mean of adjacency matrix constructed with cross-spectrum in epoch 2. The results are reported as follows: statistics value (degrees of freedom), p-value of the test, Cohen’s d effect size (or nonparametric alternative), number of epochs where significant differences were observed (E), difference estimate *µ* with 95% confidence interval. Reliable differences (significant in all three epochs) are highlighted with bold text.

**TABLE C.XII:**
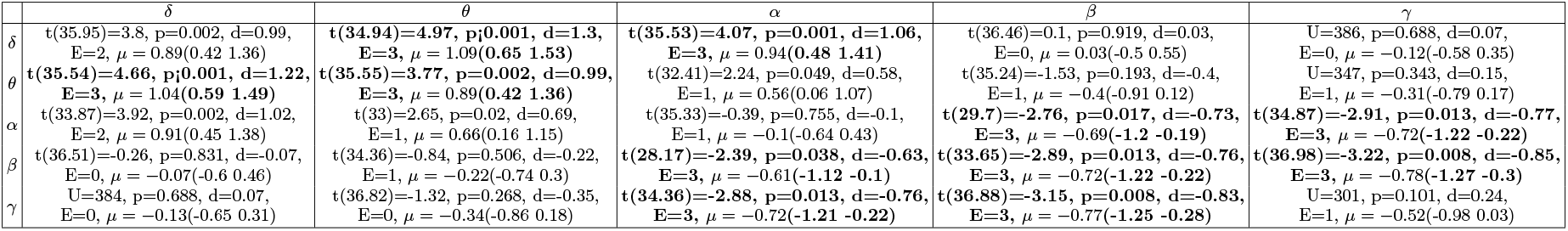
Comparisons of the mean of adjacency matrix constructed with cross-bispectrum in epoch 2. The results are reported as follows: statistics value (degrees of freedom), p-value of the test, Cohen’s d effect size (or nonparametric alternative), number of epochs where significant differences were observed (E), difference estimate *µ* with 95% confidence interval. Reliable differences (significant in all three epochs) are highlighted with bold text.

**TABLE C.XIII:**
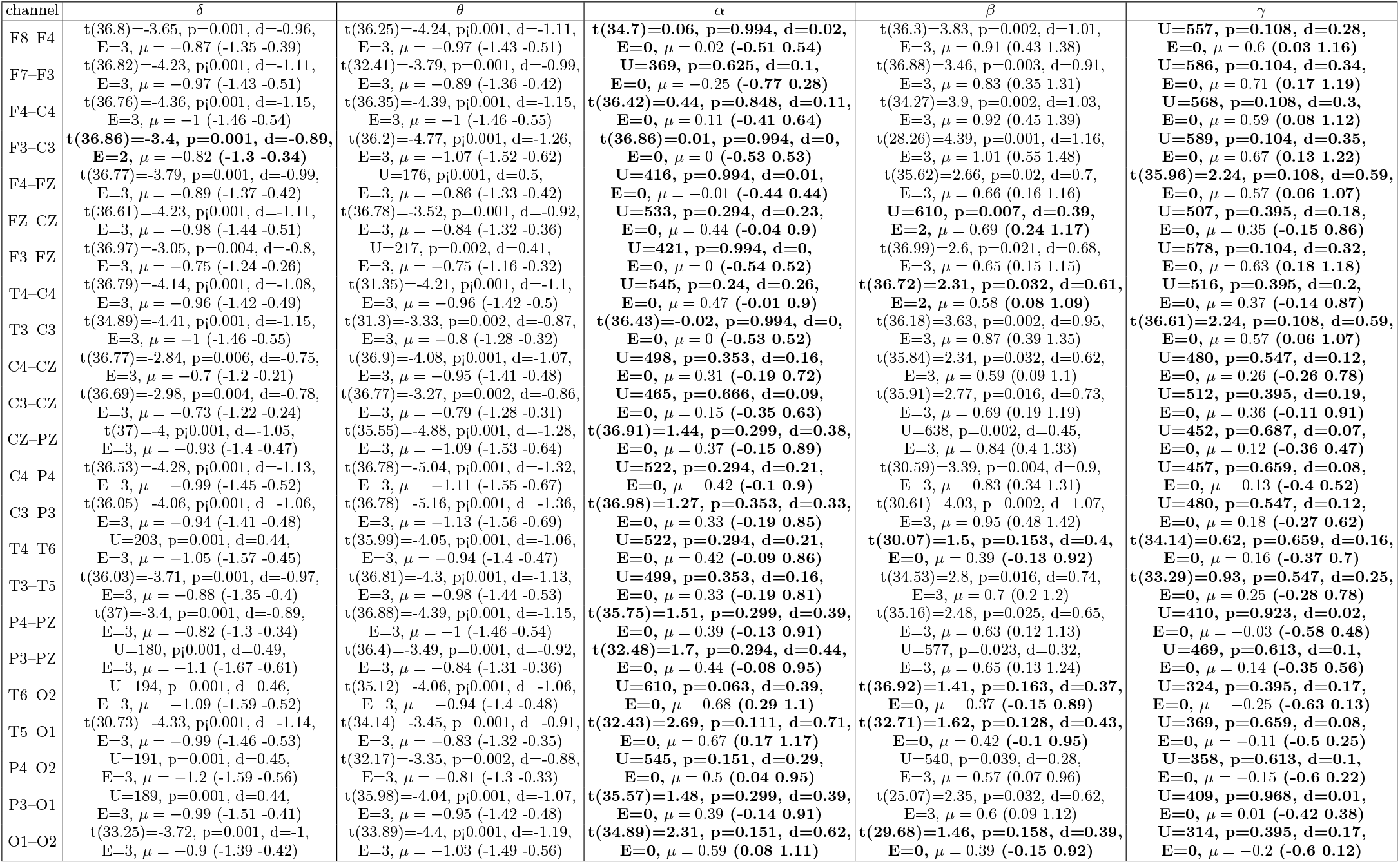
Comparisons of node strength measured with cross-spectrum in epoch 2. The results are reported as follows: statistics value (degrees of freedom), p-value of the test, Cohen’s d effect size (or nonparametric alternative), number of epochs where significant differences were observed (E), and difference estimate *µ* with 95% confidence interval (CI). Reliable differences (significant in all three epochs) are highlighted with bold text.

**TABLE C.XIV:**
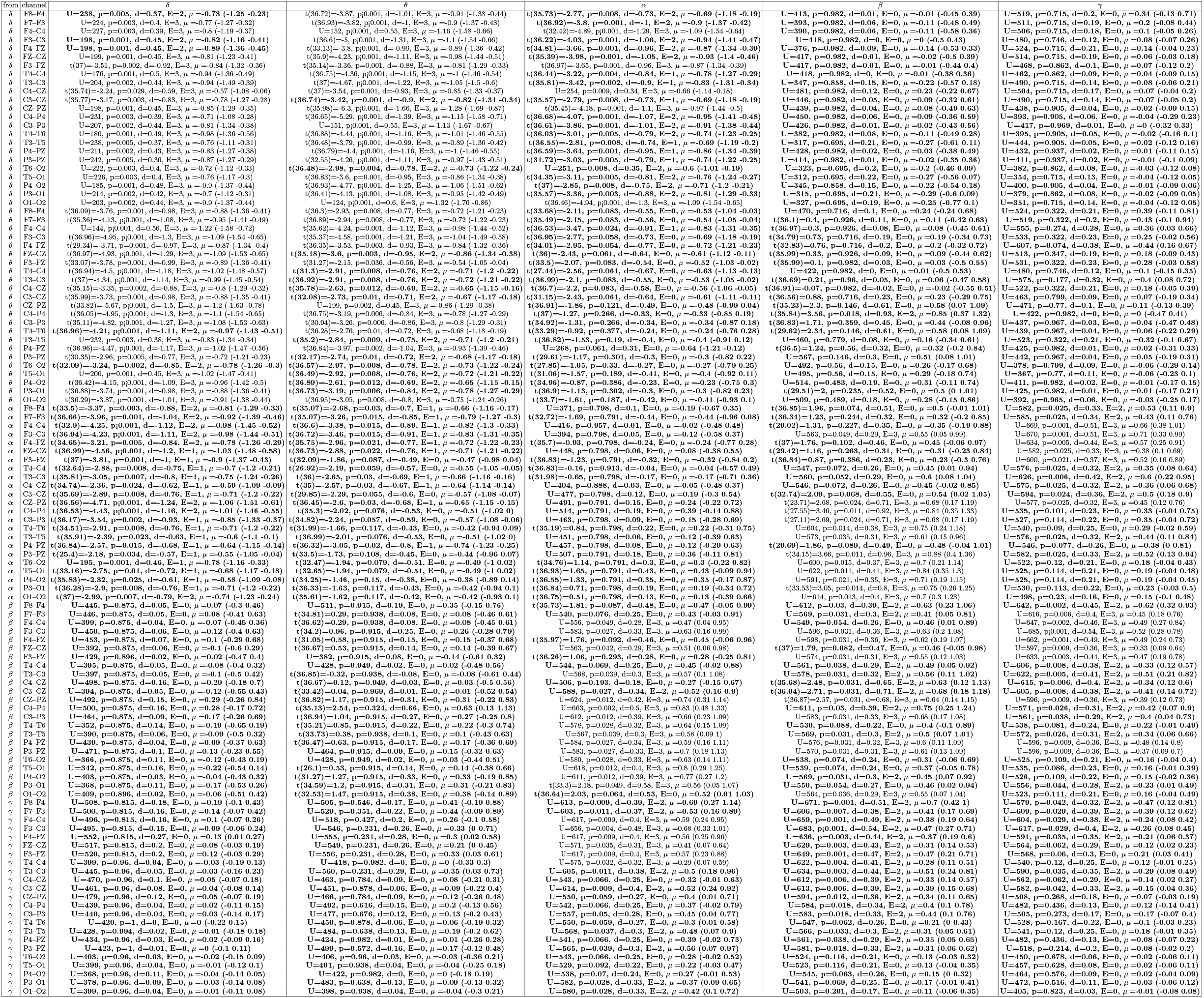
Comparisons of node strength measured with cross-bispectrum in epoch 2. The results are reported as follows: statistics value (degrees of freedom), p-value of the test, Cohen’s d effect size (or nonparametric alternative), number of epochs where significant differences were observed (E), and difference estimate *µ* with 95% confidence interval (CI). Reliable differences (significant in all three epochs) are highlighted with bold text.

**TABLE C.XV:**
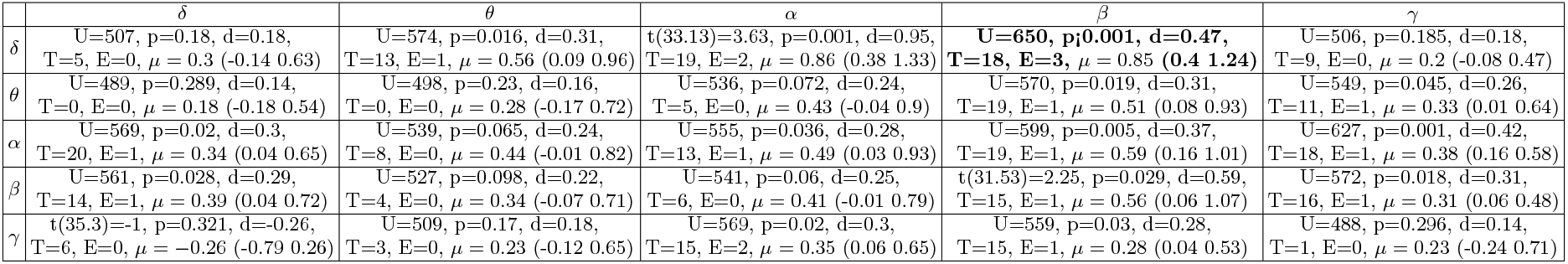
Results from epoch 2 comparing unweighted edge betweenness. The results are reported as follows: statistics value (degrees of freedom), p-value of the test, Cohen’s d effect size or nonparameteric alternative, number of thresholds where significant differences were observed (T), number of epochs where significant differences were observed (E), and group difference estimate (95% CI). Reliable differences (significant in all three epochs) are highlighted with bold text.

**TABLE C.XVI:** Results from epoch 2 comparing unweighted global vulnerability. The results are reported as follows: statistics value (degrees of freedom), p-value of the test, Cohen’s d effect size or nonparameteric alternative, number of thresholds where significant differences were observed (T), number of epochs where significant differences were observed (E), and group difference estimate (95% CI). Reliable differences (significant in all three epochs) are highlighted with bold text.

| | $\delta$ | $\theta$ | $\alpha$ | $\beta$ | $\gamma$ |
| --- | --- | --- | --- | --- | --- |
| $\delta$ | **t(27.5)=-4.11, p<0.001, d=-1.07,**<br>**T=19, E=3, $\mu = -0.94$ (-1.4 -0.48)** | **t(37)=-4.64, p<0.001, d=-1.22,**<br>**T=16, E=3, $\mu = -1.04$ (-1.49 -0.59)** | t(36.96)=-1.31, p=0.194, d=-0.35,<br>T=3, E=0, $\mu = -0.34$ (-0.87 0.18) | t(33.86)=-1.24, p=0.219, d=-0.33,<br>T=2, E=0, $\mu = -0.32$ (-0.84 0.2) | t(36.99)=-1.78, p=0.08, d=-0.47,<br>T=8, E=0, $\mu = -0.46$ (-0.97 0.06) |
| $\theta$ | t(35.35)=1.21, p=0.232, d=0.32,<br>T=4, E=0, $\mu = 0.32$ (-0.21 0.84) | U=315, p=0.104, d=0.21,<br>T=1, E=0, $\mu = -0.35$ (-0.87 0.08) | U=492, p=0.268, d=0.15,<br>T=0, E=0, $\mu = 0.18$ (-0.14 0.51) | U=348, p=0.268, d=0.15,<br>T=0, E=0, $\mu = -0.23$ (-0.77 0.27) | U=344, p=0.242, d=0.16,<br>T=2, E=0, $\mu = -0.22$ (-0.66 0.17) |
| $\alpha$ | U=505, p=0.19, d=0.17,<br>T=7, E=0, $\mu = 0.35$ (-0.14 0.88) | t(35.45)=-1.09, p=0.281, d=-0.29,<br>T=5, E=0, $\mu = -0.29$ (-0.81 0.24) | U=511, p=0.16, d=0.19,<br>T=5, E=0, $\mu = 0.28$ (-0.13 0.62) | t(34.49)=-0.74, p=0.461, d=-0.2,<br>T=0, E=0, $\mu = -0.2$ (-0.73 0.33) | U=386, p=0.605, d=0.07,<br>T=0, E=0, $\mu = -0.13$ (-0.63 0.35) |
| $\beta$ | t(30.07)=-1.04, p=0.301, d=-0.27,<br>T=6, E=0, $\mu = -0.27$ (-0.79 0.25) | t(36.97)=-0.74, p=0.462, d=-0.19,<br>T=0, E=0, $\mu = -0.19$ (-0.72 0.33) | U=453, p=0.616, d=0.07,<br>T=1, E=0, $\mu = 0.03$ (-0.07 0.12) | t(34.58)=2.73, p=0.008, d=0.72,<br>T=10, E=1, $\mu = 0.68$ (0.18 1.19) | t(37)=-2.34, p=0.023, d=-0.61,<br>T=12, E=2, $\mu = -0.59$ (-1.1 -0.08) |
| $\gamma$ | U=370, p=0.444, d=0.1,<br>T=6, E=0, $\mu = -0.12$ (-0.51 0.21) | t(36.76)=0.65, p=0.516, d=0.17,<br>T=0, E=0, $\mu = 0.17$ (-0.36 0.7) | U=456, p=0.583, d=0.07,<br>T=1, E=0, $\mu = 0.13$ (-0.34 0.6) | **t(34.2)=-3.94, p<0.001, d=-1.04,**<br>**T=14, E=3, $\mu = -0.93$ (-1.4 -0.45)** | t(36.54)=-1.63, p=0.109, d=-0.43,<br>T=7, E=0, $\mu = -0.42$ (-0.94 0.1) |

**TABLE C.XVII:**
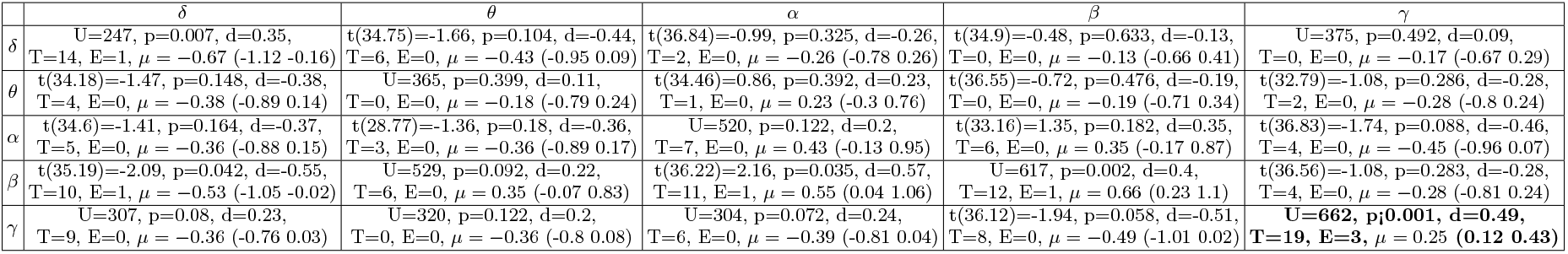
Results from epoch 2 comparing unweighted local vulnerability. The results are reported as follows: statistics value (degrees of freedom), p-value of the test, Cohen’s d effect size or nonparameteric alternative, number of thresholds where significant differences were observed (T), number of epochs where significant differences were observed (E), and group difference estimate (95% CI). Reliable differences (significant in all three epochs) are highlighted with bold text.

**TABLE C.XVIII:** Results from epoch 2 comparing weighted edge betweenness. The results are reported as follows: statistics value (degrees of freedom), p-value of the test, Cohen’s d effect size or nonparameteric alternative, number of thresholds where significant differences were observed (T), number of epochs where significant differences were observed (E), and group difference estimate (95% CI). Reliable differences (significant in all three epochs) are highlighted with bold text.

| | $\delta$ | $\theta$ | $\alpha$ | $\beta$ | $\gamma$ |
| --- | --- | --- | --- | --- | --- |
| $\delta$ | **U=608, p<0.001, d=0.47,**<br>**T=20, E=3, $\mu = 0$ (0 0.01)** | **U=618, p<0.001, d=0.49,**<br>**T=14, E=3, $\mu = 0.03$ (0 0.14)** | **U=681.5, p<0.001, d=0.55,**<br>**T=19, E=3, $\mu = 0.45$ (0.21 0.91)** | **U=671, p<0.001, d=0.51,**<br>**T=18, E=3, $\mu = 0.25$ (0.14 0.55)** | **U=639, p<0.001, d=0.45,**<br>**T=19, E=3, $\mu = 0.95$ (0.38 1.38)** |
| $\theta$ | **U=683.5, p<0.001, d=0.54,**<br>**T=10, E=3, $\mu = 0.56$ (0.3 1)** | **U=556.5, p=0.003, d=0.39,**<br>**T=10, E=3, $\mu = 0$ (0 0)** | U=448, p=0.176, d=0.18,<br>T=4, E=0, $\mu = 0$ (0 0) | U=478.5, p=0.367, d=0.12,<br>T=1, E=0, $\mu = 0.07$ (-0.02 0.24) | U=480, p=0.357, d=0.12,<br>T=1, E=0, $\mu = 0.13$ (-0.2 0.38) |
| $\alpha$ | U=504.5, p=0.065, d=0.24,<br>T=10, E=2, $\mu = 0$ (0 0) | U=461, p=0.265, d=0.15,<br>T=1, E=0, $\mu = 0$ (0 0) | U=458, p=0.354, d=0.12,<br>T=0, E=0, $\mu = 0$ (0 0) | U=442, p=0.738, d=0.04,<br>T=2, E=0, $\mu = 0.03$ (-0.15 0.27) | U=350, p=0.282, d=0.14,<br>T=1, E=0, $\mu = -0.14$ (-0.44 0.15) |
| $\beta$ | **U=686, p<0.001, d=0.54,**<br>**T=19, E=3, $\mu = 0.71$ (0.36 1.13)** | t(36.24)=0.53, p=0.6, d=0.14,<br>T=0, E=0, $\mu = 0.14$ (-0.39 0.67) | U=389.5, p=0.51, d=0.09,<br>T=0, E=0, $\mu = 0$ (0 0) | U=455, p=0.589, d=0.07,<br>T=0, E=0, $\mu = 0$ (-0.02 0.04) | **U=176, p<0.001, d=0.5,**<br>**T=18, E=3, $\mu = -0.77$ (-1.13 -0.28)** |
| $\gamma$ | **U=628, p=0.001, d=0.42,**<br>**T=18, E=3, $\mu = 1.05$ (0.45 1.45)** | U=354, p=0.311, d=0.13,<br>T=6, E=0, $\mu = -0.23$ (-0.7 0.18) | U=381, p=0.552, d=0.08,<br>T=1, E=0, $\mu = -0.05$ (-0.22 0.12) | **U=207, p=0.001, d=0.44,**<br>**T=14, E=3, $\mu = -0.65$ (-1.04 -0.27)** | **t(32)=-3.69, p=0.001, d=-0.96,**<br>**T=17, E=3, $\mu = -0.87$ (-1.34 -0.4)** |

**TABLE C.XIX:** Results from epoch 2 comparing weighted global vulnerability. The results are reported as follows: statistics value (degrees of freedom), p-value of the test, Cohen’s d effect size or nonparameteric alternative, number of thresholds where significant differences were observed (T), number of epochs where significant diffferences were observed (E), group difference estimate (95% CI). Reliable differences (significant in all three epochs) are highlighted with bold text.

| | $\delta$ | $\theta$ | $\alpha$ | $\beta$ | $\gamma$ |
| --- | --- | --- | --- | --- | --- |
| $\delta$ | U=389, p=0.638, d=0.06,<br>T=3, E=0, $\mu = 0$ (0 0) | U=387, p=0.616, d=0.07,<br>T=2, E=0, $\mu = 0$ (0 0) | U=467, p=0.472, d=0.1,<br>T=3, E=0, $\mu = 0$ (0 0) | **U=645, p<0.001, d=0.46,**<br>**T=19, E=3, $\mu = 0.19$ (0.06 0.48)** | t(36.04)=1.44, p=0.154, d=0.38,<br>T=2, E=0, $\mu = 0.37$ (-0.14 0.89) |
| $\theta$ | U=377, p=0.511, d=0.09,<br>T=3, E=0, $\mu = 0$ (0 0) | U=483, p=0.333, d=0.13,<br>T=0, E=0, $\mu = 0$ (0 0.17) | U=388, p=0.627, d=0.07,<br>T=0, E=0, $\mu = 0$ (0 0) | U=386, p=0.605, d=0.07,<br>T=0, E=0, $\mu = -0.01$ (-0.06 0.03) | U=475, p=0.399, d=0.11,<br>T=0, E=0, $\mu = 0.22$ (-0.31 0.74) |
| $\alpha$ | U=483, p=0.333, d=0.13,<br>T=5, E=0, $\mu = 0$ (0 0) | U=388, p=0.627, d=0.07,<br>T=0, E=0, $\mu = 0$ (0 0) | U=400, p=0.763, d=0.04,<br>T=0, E=0, $\mu = 0$ (0 0) | U=388, p=0.627, d=0.07,<br>T=0, E=0, $\mu = 0$ (0 0) | U=355, p=0.318, d=0.13,<br>T=2, E=0, $\mu = -0.22$ (-0.74 0.3) |
| $\beta$ | **U=644, p<0.001, d=0.46,**<br>**T=10, E=3, $\mu = 0.4$ (0.03 0.62)** | U=379, p=0.531, d=0.08,<br>T=0, E=0, $\mu = 0$ (0 0) | U=459, p=0.552, d=0.08,<br>T=0, E=0, $\mu = 0$ (0 0) | U=458, p=0.562, d=0.08,<br>T=2, E=0, $\mu = 0$ (0 0) | **U=176, p<0.001, d=0.5,**<br>**T=18, E=3, $\mu = -0.64$ (-0.96 -0.3)** |
| $\gamma$ | U=532, p=0.083, d=0.23,<br>T=4, E=0, $\mu = 0.35$ (-0.03 0.83) | U=446, p=0.694, d=0.05,<br>T=0, E=0, $\mu = 0.13$ (-0.28 0.6) | U=396, p=0.717, d=0.05,<br>T=0, E=0, $\mu = -0.08$ (-0.51 0.28) | **t(35.65)=-3.68, p=0.001, d=-0.96,**<br>**T=20, E=3, $\mu = -0.87$ (-1.34 -0.4)** | **U=185, p<0.001, d=0.48,**<br>**T=18, E=3, $\mu = -0.96$ (-1.5 -0.38)** |

**TABLE C.XX:** Results from epoch 2 comparing weighted local vulnerability. The results are reported as follows: statistics value (degrees of freedom), p-value of the test, Cohen’s d effect size or nonparameteric alternative, number of thresholds where significant differences were observed (T), number of epochs where significant differences were observed (E), and group difference estimate (95% CI). Reliable differences (significant in all three epochs) are highlighted with bold text.

| | $\delta$ | $\theta$ | $\alpha$ | $\beta$ | $\gamma$ |
| --- | --- | --- | --- | --- | --- |
| $\delta$ | U=644, p\|0.001, d=0.46,<br>T=17, E=3, $\mu = 0.79$ (0.36 1.22) | U=657, p\|0.001, d=0.48,<br>T=18, E=3, $\mu = 0.86$ (0.46 1.29) | U=213, p=0.001, d=0.42,<br>T=14, E=3, $\mu = -0.76$ (-1.13 -0.39) | U=176, p\|0.001, d=0.5,<br>T=14, E=3, $\mu = -0.7$ (-0.97 -0.4) | U=456, p=0.583, d=0.07,<br>T=0, E=0, $\mu = 0.1$ (-0.28 0.47) |
| $\theta$ | t(28.75)=3.75, p\|0.001, d=0.98,<br>T=19, E=3, $\mu = 0.88$ (0.41 1.35) | t(36.76)=3.51, p=0.001, d=0.92,<br>T=15, E=3, $\mu = 0.84$ (0.36 1.32) | U=263, p=0.014, d=0.32,<br>T=12, E=2, $\mu = -0.57$ (-0.91 -0.13) | t(33.55)=-5.08, p\|0.001, d=-1.34,<br>T=16, E=3, $\mu = -1.12$ (-1.56 -0.68) | U=349, p=0.275, d=0.15,<br>T=0, E=0, $\mu = -0.15$ (-0.46 0.13) |
| $\alpha$ | t(36.95)=3.9, p\|0.001, d=1.02,<br>T=19, E=3, $\mu = 0.92$ (0.45 1.38) | t(34.18)=3.25, p=0.002, d=0.86,<br>T=14, E=3, $\mu = 0.79$ (0.3 1.28) | U=212, p=0.001, d=0.42,<br>T=14, E=3, $\mu = -0.82$ (-1.29 -0.34) | t(28.29)=-5.33, p\|0.001, d=-1.41,<br>T=15, E=3, $\mu = -1.16$ (-1.6 -0.72) | U=349, p=0.275, d=0.15,<br>T=5, E=0, $\mu = -0.21$ (-0.62 0.14) |
| $\beta$ | t(33.58)=3.73, p\|0.001, d=0.97,<br>T=18, E=3, $\mu = 0.88$ (0.41 1.35) | U=610, p=0.003, d=0.39,<br>T=11, E=3, $\mu = 0.7$ (0.24 1.19) | U=271, p=0.02, d=0.3,<br>T=12, E=1, $\mu = -0.5$ (-0.84 -0.07) | U=177, p\|0.001, d=0.5,<br>T=14, E=3, $\mu = -0.87$ (-1.24 -0.51) | U=320, p=0.122, d=0.2,<br>T=6, E=0, $\mu = -0.26$ (-0.59 0.05) |
| $\gamma$ | U=547, p=0.048, d=0.26,<br>T=10, E=1, $\mu = 0.48$ (0 1.06) | t(32.94)=1.24, p=0.222, d=0.33,<br>T=4, E=0, $\mu = 0.33$ (-0.2 0.85) | t(33.97)=-0.65, p=0.521, d=-0.17,<br>T=0, E=0, $\mu = -0.17$ (-0.69 0.36) | t(32.39)=-3.72, p\|0.001, d=-0.98,<br>T=16, E=3, $\mu = -0.89$ (-1.37 -0.41) | t(35.36)=-3.19, p=0.002, d=-0.83,<br>T=11, E=3, $\mu = -0.77$ (-1.26 -0.29) |

## Appendix D: Results for epoch 3

In this section, we only report the results of the statistical comparisons and corresponding figures using data from the third epoch.

The comparisons of average coupling computed with cross-spectrum and cross-bispectrum are shown in Tables D.XXI and D.XXII, respectively. Additionally, the corresponding figures of connectivity matrices computed with cross-spectrum and cross-bispectrum are shown in Figures D.12 and D.13.

The comparisons of node strength computed with cross-spectrum and cross-bispectrum are shown in Table D.XXIII and Figure D.14, and Table D.XXIV and Figure D.15, respectively.

Results of comparisons of the unweighted multilayer network metrics are reported in Tables D.XXV, D.XXVI and D.XXVII for edge betweenness, global vulnerability and local vulnerability, respectively. The corresponding figures are; edge betweenness (Figure D.16), global vulnerability (Figure D.17) and local vulnerability (Figure D.18).

Results of comparisons of the weighted multilayer network metrics are reported in Tables D.XXVIII, D.XXIX and D.XXX for edge betweenness, global vulnerability and local vulnerability, respectively. The corresponding figures are; edge betweenness (Figure D.19), global vulnerability (Figure D.20) and local vulnerability (Figure D.21).

**FIG. D.12:**
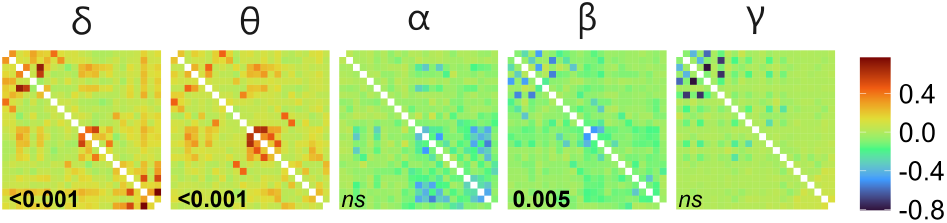
Difference of average connectivity matrices (*AD − HC*) measured with cross-spectrum in epoch 3. For visualisation purposes, the values were min-max normalised. Digits in black denote a *p*-value testing for the difference in global coupling (*p <* 0.05 in bold, in italics otherwise).

**TABLE D.XXI:** Comparisons of the mean of adjacency matrix constructed with cross-spectrum in epoch 3. The results are reported as follows: statistics value (degrees of freedom), p-value of the test, Cohen’s d effect size (or nonparametric alternative), number of epochs where significant differences were observed (E), difference estimate *µ* with 95% confidence interval. Reliable differences (significant in all three epochs) are highlighted with bold text.

| Frequency band | Test | Difference estimate (95% CI) |
| --- | --- | --- |
| $\alpha$ | $t(34.87)=-2.12, p=0.049, d=-0.55, E=1$ | $\mu = -0.54 (-1.04 -0.03)$ |
| $\beta$ | $U=231, p=0.005, d=0.39, E=3$ | $\mu = -0.72 (-1.19 -0.2)$ |
| $\delta$ | **$t(34.62)=4.82, p<0.001, d=1.26, E=3$** | $\mu = 1.07 (0.62 1.51)$ |
| $\gamma$ | $U=355, p=0.318, d=0.13, E=0$ | $\mu = -0.27 (-0.85 0.31)$ |
| $\theta$ | **$t(32.41)=4.93, p<0.001, d=1.3, E=3$** | $\mu = 1.1 (0.65 1.54)$ |

**FIG. D.13:**
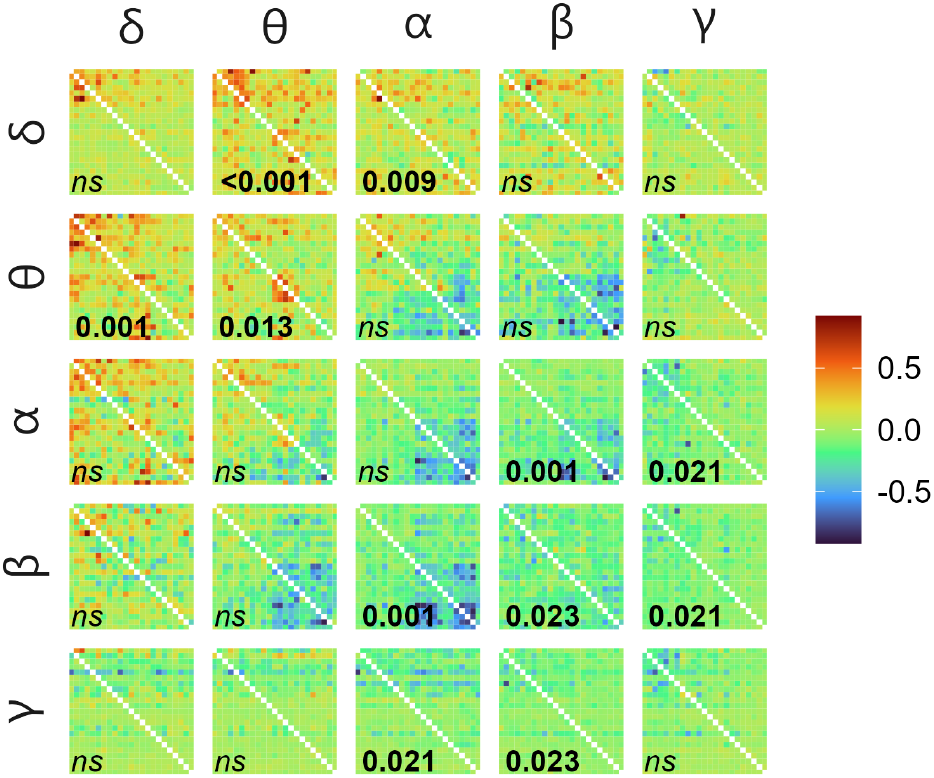
Difference of average connectivity matrices (*AD − HC*) measured with cross-bispectrum in epoch 3 with input frequency on the vertical facets and output frequency on the horizontal. For visualisation purposes, the values were min-max normalised. Digits in white denote a *p*-value testing for the difference in global coupling (*p <* 0.05 in bold, in italics otherwise).

**TABLE D.XXII:**
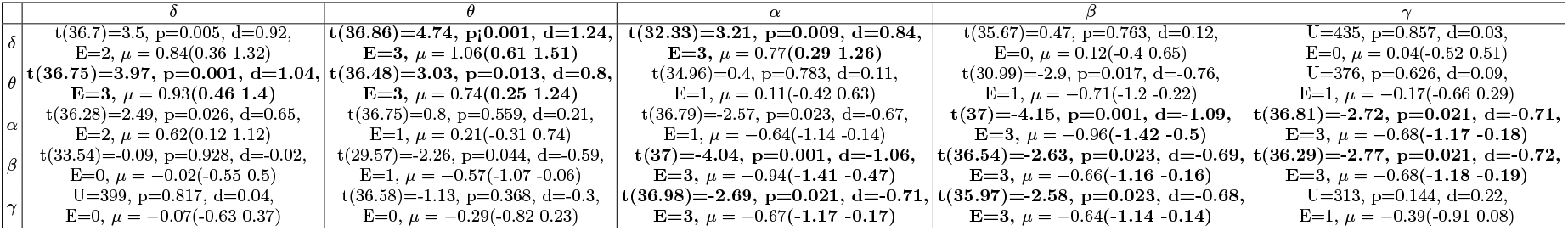
Comparisons of the mean of adjacency matrix constructed with cross-bispectrum in epoch 3. The results are reported as follows: statistics value (degrees of freedom), p-value of the test, Cohen’s d effect size (or nonparametric alternative), number of epochs where significant differences were observed (E), difference estimate *µ* with 95% confidence interval. Reliable differences (significant in all three epochs) are highlighted with bold text.

**FIG. D.14:**
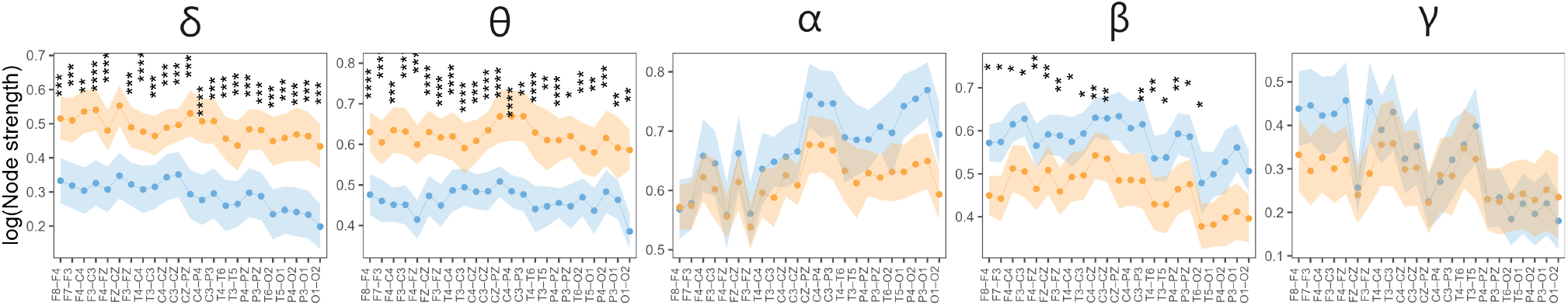
Node strength (min-max normalised) measured with CS in epoch 3 of HC (blue) and AD (orange): mean with 95% confidence intervals. Significant differences (*p ≤* 0.05) observed in at least ten thresholded networks are encoded by asterisks. The number of asterisks corresponds to the p-value (FDR corrected), i.e. *p ≤* 0.0001 “****”, *p ≤* 0.001 “***”, *p ≤* 0.01 “**”, and *p ≤* 0.05 “*”.

**TABLE D.XXIII:**
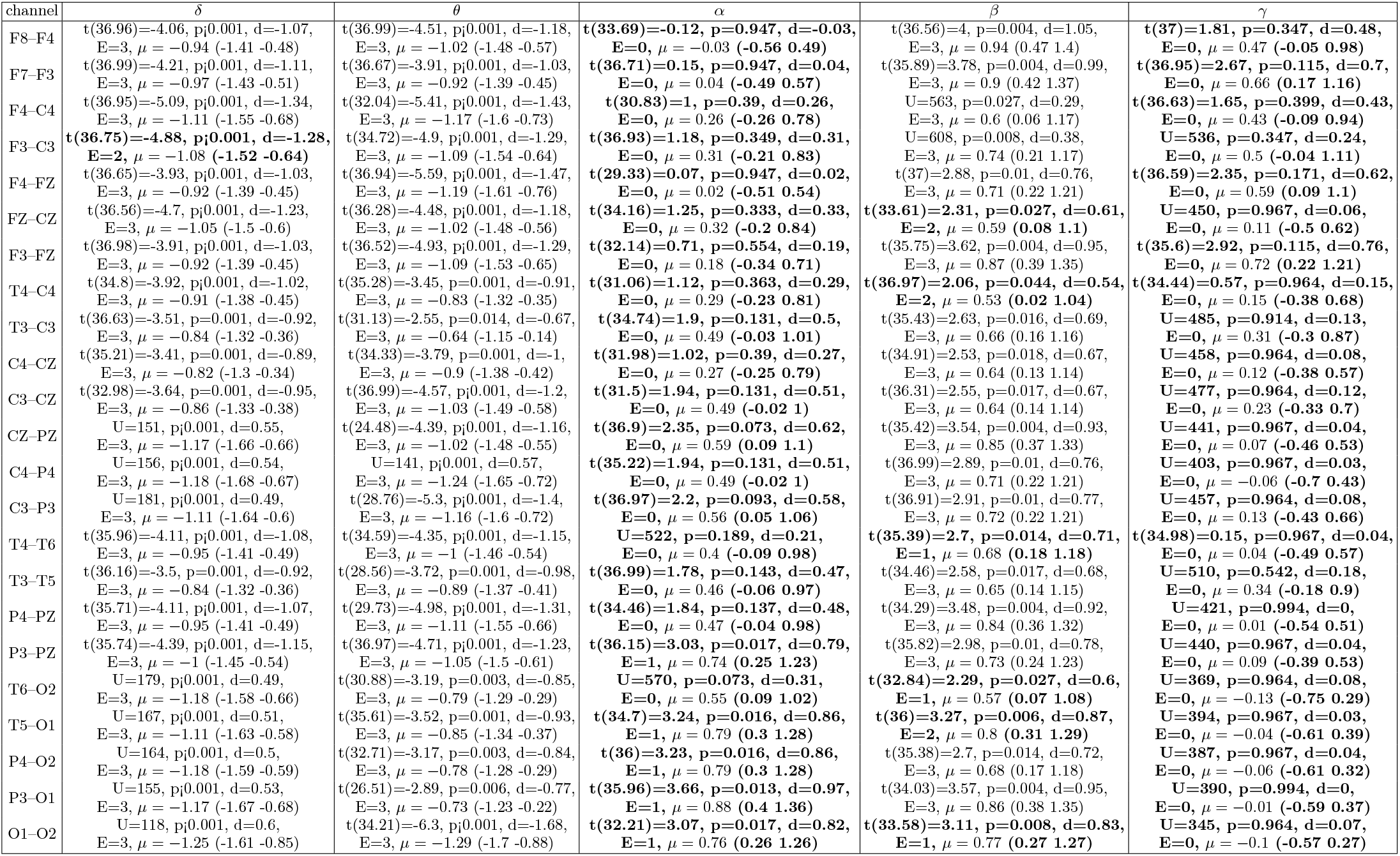
Comparisons of node strength measured with cross-spectrum in epoch 3. The results are reported as follows: statistics value (degrees of freedom), p-value of the test, Cohen’s d effect size (or nonparametric alternative), number of epochs where significant differences were observed (E), and difference estimate *µ* with 95% confidence interval (CI). Reliable differences (significant in all three epochs) are highlighted with bold text.

**FIG. D.15:**
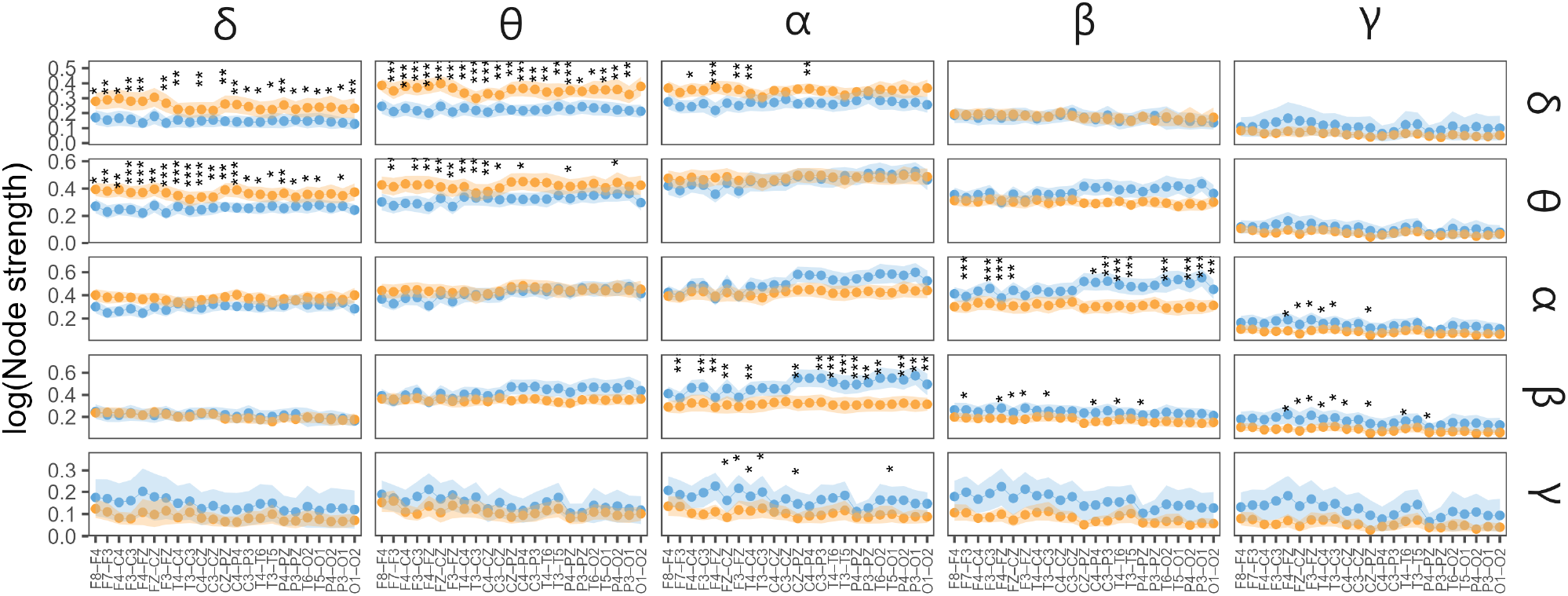
Node strength (min-max normalised) measured with CBS in epoch 3 of HC (blue) and AD (orange): mean with 95% confidence intervals. The input frequency is on the vertical facets, and the output frequency is on the horizontal. Significant differences (*p ≤* 0.05) observed in at least ten thresholded networks are encoded by asterisks. The number of asterisks corresponds to the p-value (FDR corrected), i.e. *p ≤* 0.0001 “****”, *p ≤* 0.001 “***”, *p ≤* 0.01 “**”, and *p ≤* 0.05 “*”.

**TABLE D.XXIV:**
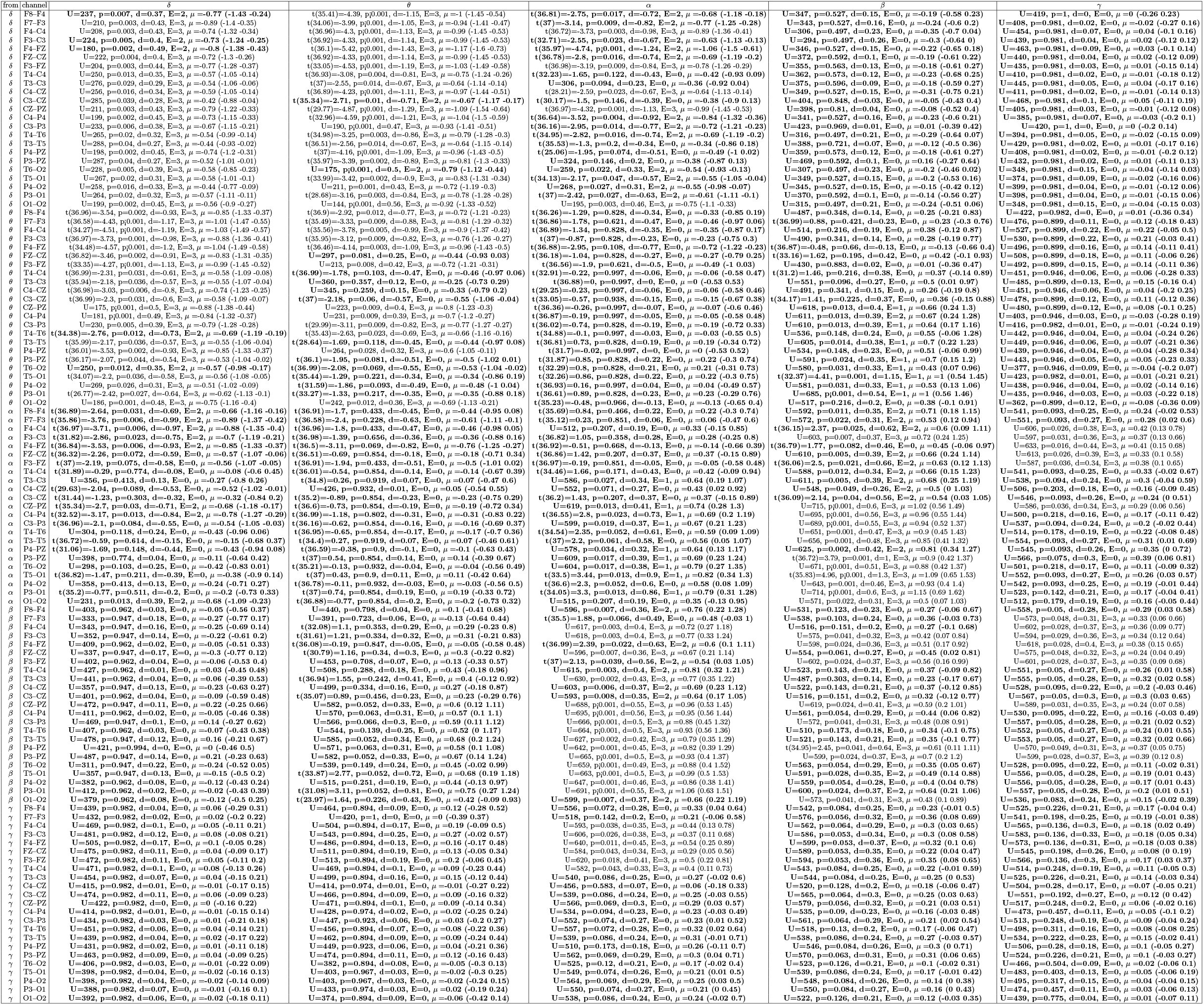
Comparisons of node strength measured with cross-bispectrum in epoch 3. The results are reported as follows: statistics value (degrees of freedom), p-value of the test, Cohen’s d effect size (or nonparametric alternative), number of epochs where significant differences were observed (E), and difference estimate *µ* with 95% confidence interval (CI). Reliable differences (significant in all three epochs) are highlighted with bold text.

**TABLE D.XXV:**
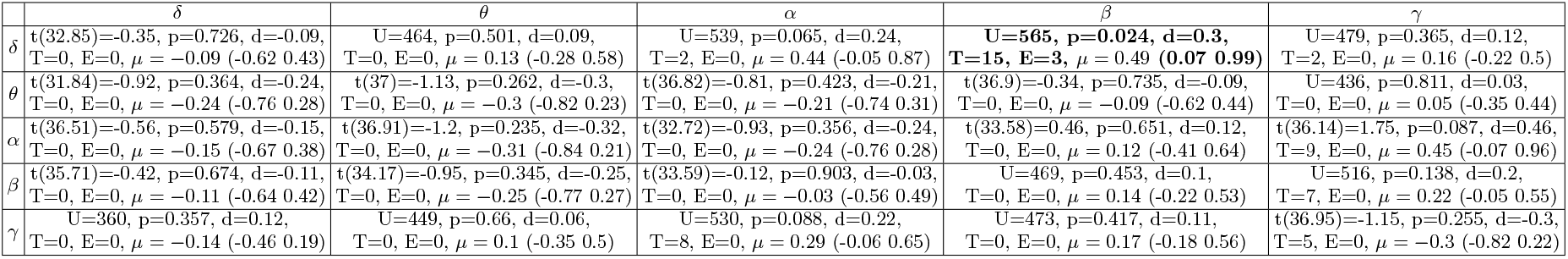
Results from epoch 3 comparing unweighted edge betweenness. The results are reported as follows: statistics value (degrees of freedom), p-value of the test, Cohen’s d effect size or nonparameteric alternative, number of thresholds where significant differences were observed (T), number of epochs where significant differences were observed (E), group difference estimate (95% CI). Reliable differences (significant in all three epochs) are highlighted with bold text.

**TABLE D.XXVI:**
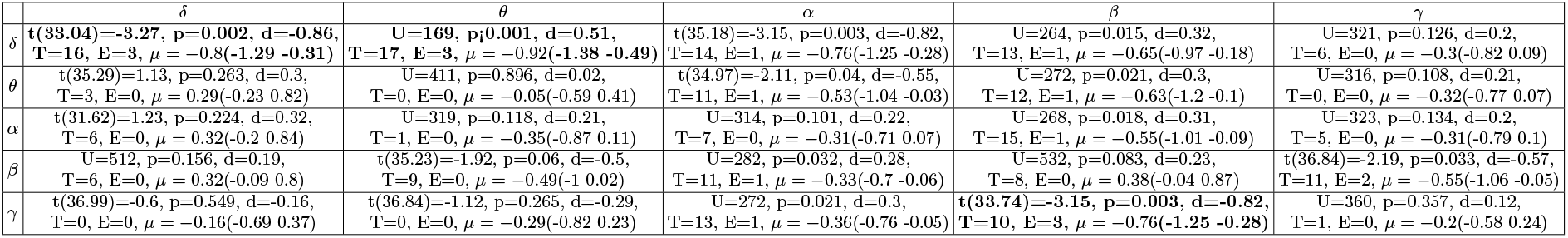
Results from epoch 3 comparing unweighted global vulnerability. The results are reported as follows: statistics value (degrees of freedom), p-value of the test, Cohen’s d effect size or nonparameteric alternative, number of thresholds where significant differences were observed (T), number of epochs where significant differences were observed (E), group difference estimate (95% CI). Reliable differences (significant in all three epochs) are highlighted with bold text.

**TABLE D.XXVII:**
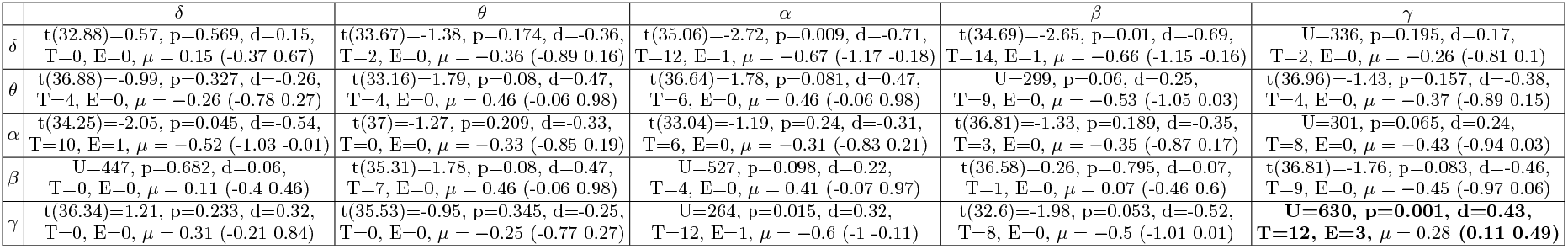
Results from epoch 3 comparing unweighted local vulnerability. The results are reported as follows: statistics value (degrees of freedom), p-value of the test, Cohen’s d effect size or nonparameteric alternative, number of thresholds where significant differences were observed (T), number of epochs where significant differences were observed (E), group difference estimate (95% CI). Reliable differences (significant in all three epochs) are highlighted with bold text.

| | $\delta$ | $\theta$ | $\alpha$ | $\beta$ | $\gamma$ |
| --- | --- | --- | --- | --- | --- |
| $\delta$ | t(32.88)=0.57, p=0.569, d=0.15, T=0, E=0, $\mu = 0.15$ (-0.37 0.67) | t(33.67)=-1.38, p=0.174, d=-0.36, T=2, E=0, $\mu = -0.36$ (-0.89 0.16) | t(35.06)=-2.72, p=0.009, d=-0.71, T=12, E=1, $\mu = -0.67$ (-1.17 -0.18) | t(34.69)=-2.65, p=0.01, d=-0.69, T=14, E=1, $\mu = -0.66$ (-1.15 -0.16) | U=336, p=0.195, d=0.17, T=2, E=0, $\mu = -0.26$ (-0.81 0.1) |
| $\theta$ | t(36.88)=-0.99, p=0.327, d=-0.26, T=4, E=0, $\mu = -0.26$ (-0.78 0.27) | t(33.16)=1.79, p=0.08, d=0.47, T=4, E=0, $\mu = 0.46$ (-0.06 0.98) | t(36.64)=1.78, p=0.081, d=0.47, T=6, E=0, $\mu = 0.46$ (-0.06 0.98) | U=299, p=0.06, d=0.25, T=9, E=0, $\mu = -0.53$ (-1.05 0.03) | t(36.96)=-1.43, p=0.157, d=-0.38, T=4, E=0, $\mu = -0.37$ (-0.89 0.15) |
| $\alpha$ | t(34.25)=-2.05, p=0.045, d=-0.54, T=10, E=1, $\mu = -0.52$ (-1.03 -0.01) | t(37)=-1.27, p=0.209, d=-0.33, T=0, E=0, $\mu = -0.33$ (-0.85 0.19) | t(33.04)=-1.19, p=0.24, d=-0.31, T=6, E=0, $\mu = -0.31$ (-0.83 0.21) | t(36.81)=-1.33, p=0.189, d=-0.35, T=3, E=0, $\mu = -0.35$ (-0.87 0.17) | U=301, p=0.065, d=0.24, T=8, E=0, $\mu = -0.43$ (-0.94 0.03) |
| $\beta$ | U=447, p=0.682, d=0.06, T=0, E=0, $\mu = 0.11$ (-0.4 0.46) | t(35.31)=1.78, p=0.08, d=0.47, T=7, E=0, $\mu = 0.46$ (-0.06 0.98) | U=527, p=0.098, d=0.22, T=4, E=0, $\mu = 0.41$ (-0.07 0.97) | t(36.58)=0.26, p=0.795, d=0.07, T=1, E=0, $\mu = 0.07$ (-0.46 0.6) | t(36.81)=-1.76, p=0.083, d=-0.46, T=9, E=0, $\mu = -0.45$ (-0.97 0.06) |
| $\gamma$ | t(36.34)=1.21, p=0.233, d=0.32, T=0, E=0, $\mu = 0.31$ (-0.21 0.84) | t(35.53)=-0.95, p=0.345, d=-0.25, T=0, E=0, $\mu = -0.25$ (-0.77 0.27) | U=264, p=0.015, d=0.32, T=12, E=1, $\mu = -0.6$ (-1 -0.11) | t(32.6)=-1.98, p=0.053, d=-0.52, T=8, E=0, $\mu = -0.5$ (-1.01 0.01) | U=630, p=0.001, d=0.43, T=12, E=3, $\mu = 0.28$ (0.11 0.49) |

**TABLE D.XXVIII:**
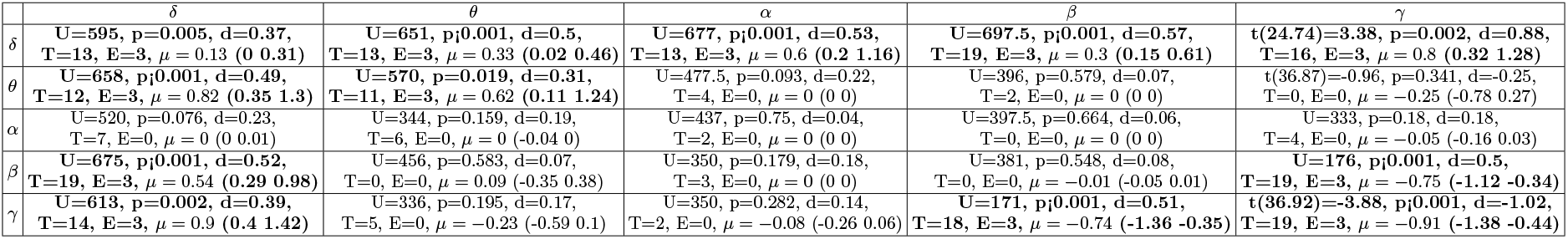
Results from epoch 3 comparing weighted edge betweenness. The results are reported as follows: statistics value (degrees of freedom), p-value of the test, Cohen’s d effect size or nonparameteric alternative, number of thresholds where significant differences were observed (T), number of epochs where significant differences were observed (E), group difference estimate (95% CI). Reliable differences (significant in all three epochs) are highlighted with bold text.

**TABLE D.XXIX:** Results from epoch 3 comparing weighted global vulnerability. The results are reported as follows: statistics value (degrees of freedom), p-value of the test, Cohen’s d effect size or nonparameteric alternative, number of thresholds where significant differences were observed (T), number of epochs where significant differences were observed (E), group difference estimate (95% CI). Reliable differences (significant in all three epochs) are highlighted with bold text.

| | $\delta$ | $\theta$ | $\alpha$ | $\beta$ | $\gamma$ |
| --- | --- | --- | --- | --- | --- |
| $\delta$ | U=466, p=0.482, d=0.09, T=1, E=0, $\mu = 0$ (0 0) | U=451, p=0.638, d=0.06, T=2, E=0, $\mu = 0$ (0 0) | U=514, p=0.147, d=0.19, T=4, E=0, $\mu = 0$ (0 0) | U=626, p=0.001, d=0.42, T=17, E=3, $\mu = 0.23$ (0.02 0.95) | U=581, p=0.012, d=0.33, T=15, E=2, $\mu = 0.68$ (0.2 1.12) |
| $\theta$ | U=453, p=0.616, d=0.07, T=2, E=0, $\mu = 0$ (0 0) | U=441, p=0.752, d=0.04, T=0, E=0, $\mu = 0$ (0 0) | U=399, p=0.752, d=0.04, T=0, E=0, $\mu = 0$ (0 0) | U=440, p=0.763, d=0.04, T=0, E=0, $\mu = 0$ (0 0) | t(32.07)=0.4, p=0.69, d=0.1, T=0, E=0, $\mu = 0.1$ (-0.42 0.63) |
| $\alpha$ | U=517, p=0.134, d=0.2, T=2, E=0, $\mu = 0$ (0 0) | U=404, p=0.811, d=0.03, T=0, E=0, $\mu = 0$ (0 0) | U=389, p=0.638, d=0.06, T=0, E=0, $\mu = 0$ (-0.12 0.01) | U=404, p=0.811, d=0.03, T=0, E=0, $\mu = 0$ (0 0) | t(30.11)=-1.11, p=0.275, d=-0.29, T=2, E=0, $\mu = -0.29$ (-0.8 0.23) |
| $\beta$ | U=659, p<0.001, d=0.49, T=10, E=3, $\mu = 0.65$ (0.03 1.1) | U=390, p=0.649, d=0.06, T=0, E=0, $\mu = 0$ (0 0) | U=451, p=0.638, d=0.06, T=0, E=0, $\mu = 0$ (0 0) | U=365, p=0.399, d=0.11, T=0, E=0, $\mu = 0$ (0 0) | U=135, p<0.001, d=0.58, T=20, E=3, $\mu = -0.95$ (-1.32 -0.55) |
| $\gamma$ | U=453, p=0.616, d=0.07, T=1, E=0, $\mu = 0.15$ (-0.41 0.7) | U=442, p=0.74, d=0.04, T=0, E=0, $\mu = 0.02$ (-0.26 0.27) | U=388, p=0.627, d=0.07, T=0, E=0, $\mu = -0.14$ (-0.62 0.4) | t(36.03)=-4.75, p<0.001, d=-1.24, T=16, E=3, $\mu = -1.06$ (-1.5 -0.61) | U=176, p<0.001, d=0.5, T=18, E=3, $\mu = -0.99$ (-1.4 -0.51) |

**TABLE D.XXX:**
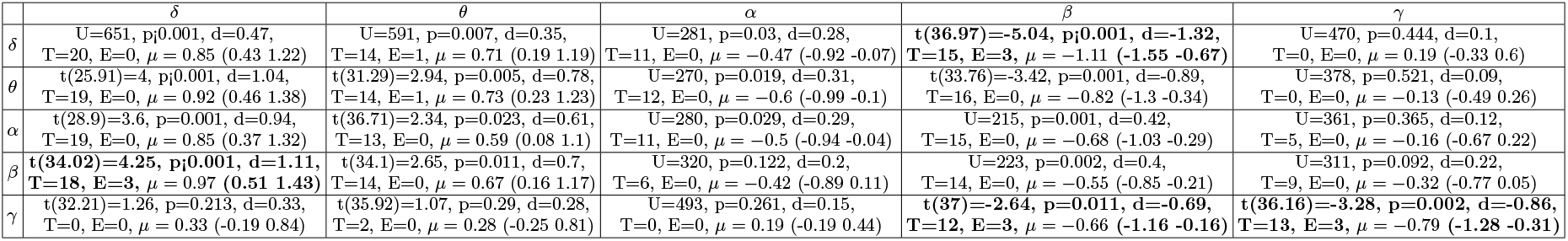
Results from epoch 3 comparing weighted local vulnerability. The results are reported as follows: statistics value (degrees of freedom), p-value of the test, Cohen’s d effect size or nonparameteric alternative, number of thresholds where significant differences were observed (T), number of epochs where significant differences were observed (E), group difference estimate (95% CI). Reliable differences (significant in all three epochs) are highlighted with bold text.

**FIG. D.16:**
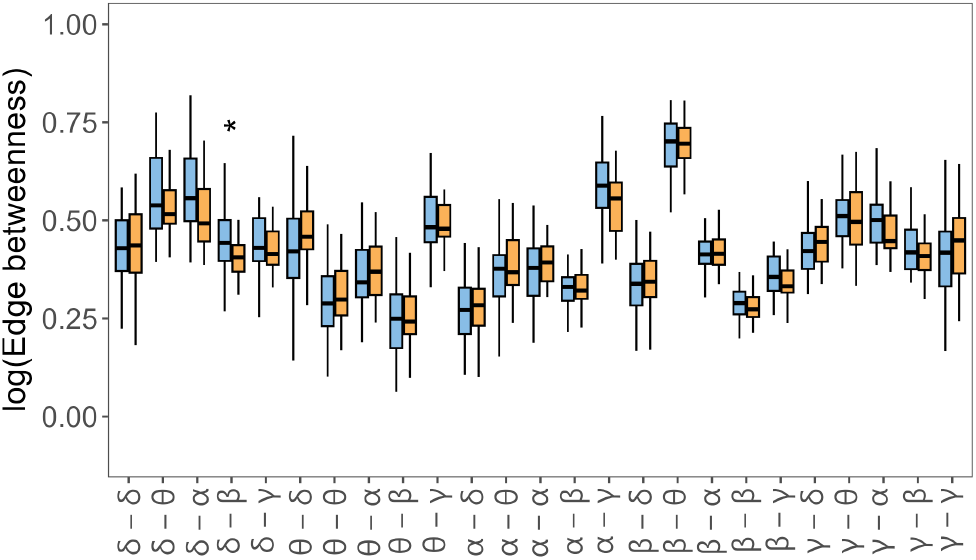
Importance of each type of frequency coupling of HC (blue) and AD (orange) measured by edge betweenness in epoch 3. Significant differences (*p ≤* 0.05) observed in at least ten thresholded networks are encoded by asterisks. The number of asterisks corresponds to the p-value (FDR corrected), i.e. *p ≤* 0.0001 “****”, *p ≤* 0.001 “***”, *p ≤* 0.01 “**”, and *p ≤* 0.05 “*”.

**FIG. D.17:**
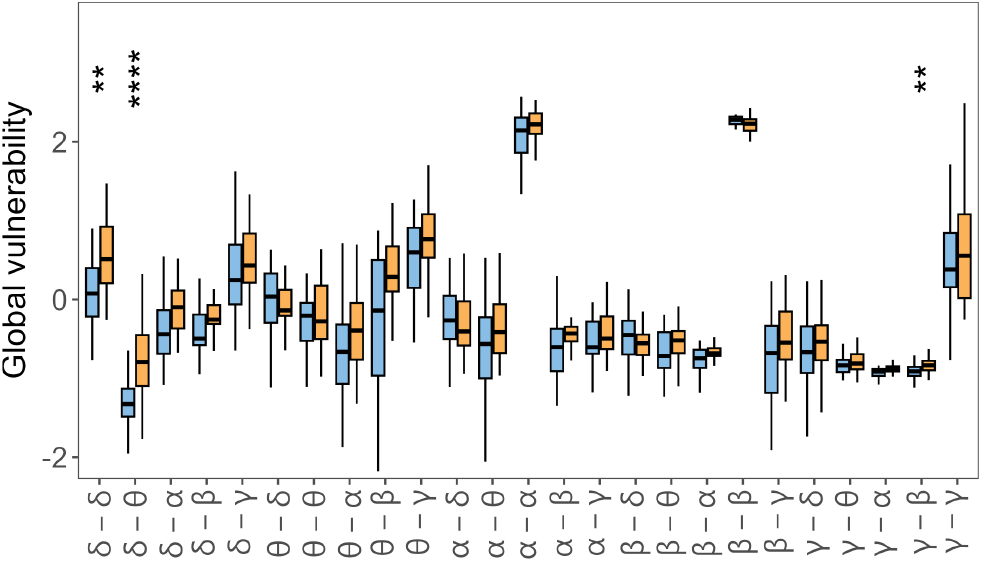
Global vulnerability of HC (blue) and AD (orange) in epoch 3. Significant differences (*p ≤* 0.05) observed in at least ten thresholded networks are encoded by asterisks. The number of asterisks corresponds to the p-value (FDR corrected), i.e. *p ≤* 0.0001 “****”, *p ≤* 0.001 “***”, *p ≤* 0.01 “**”, and *p ≤* 0.05 “*”.

**FIG. D.18:**
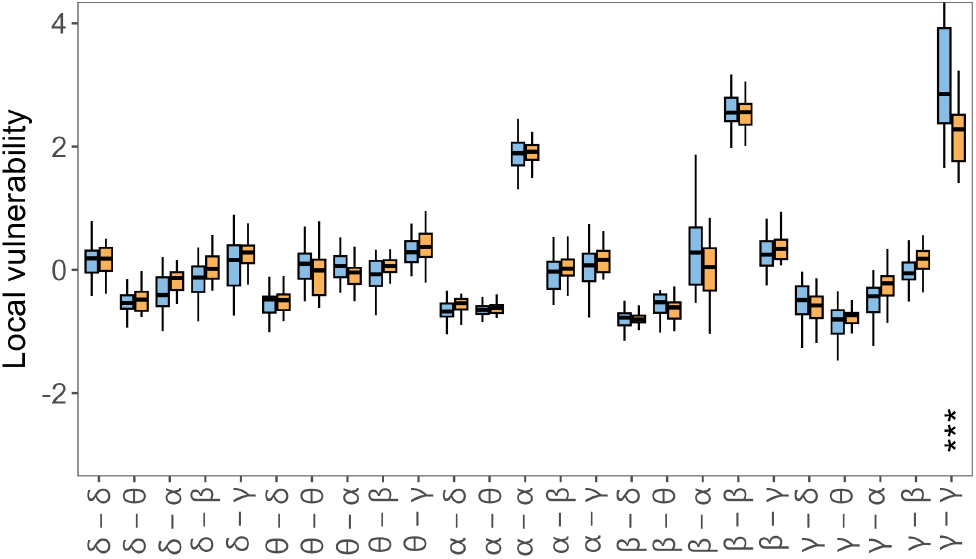
Local vulnerability of HC (blue) and AD (orange) in epoch 3. Significant differences (*p ≤* 0.05) observed in at least ten thresholded networks are encoded by asterisks. The number of asterisks corresponds to the p-value (FDR corrected), i.e. *p ≤* 0.0001 “****”, *p ≤* 0.001 “***”, *p ≤* 0.01 “**”, and *p ≤* 0.05 “*”.

**FIG. D.19:**
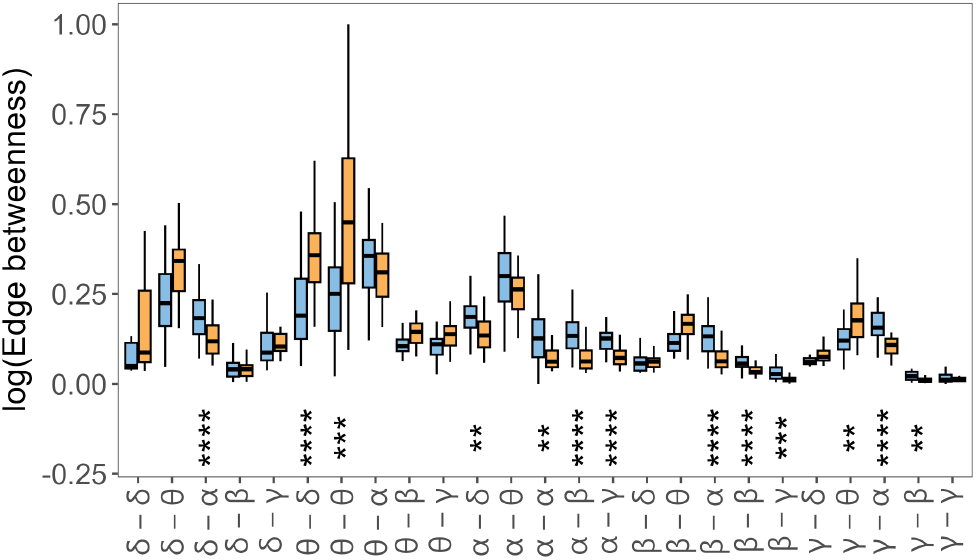
Importance of each type of frequency coupling of HC (blue) and AD (orange) measured by weighted edge betweenness in epoch 3. Significant differences (*p ≤* 0.05) observed in at least ten thresholded networks are encoded by asterisks. The number of asterisks corresponds to the p-value (FDR corrected), i.e. *p ≤* 0.0001 “****”, *p ≤* 0.001 “***”, *p ≤* 0.01 “**”, and *p ≤* 0.05 “*”.

**FIG. D.20:**
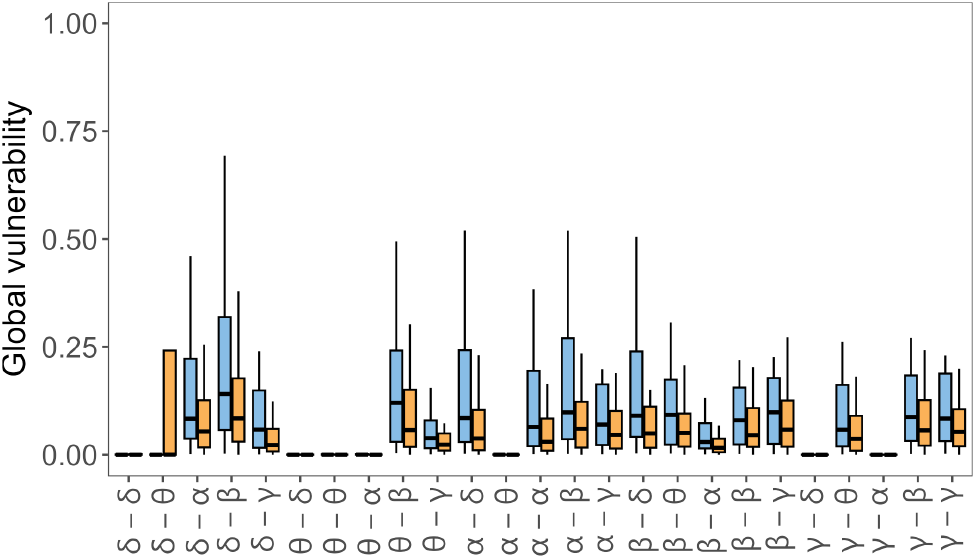
Weighted global vulnerability of HC (blue) and AD (orange) in epoch 3. Significant differences (*p ≤* 0.05) observed in at least ten thresholded networks are encoded by asterisks. The number of asterisks corresponds to the p-value (FDR corrected), i.e. *p ≤* 0.0001 “****”, *p ≤* 0.001 “***”, *p ≤* 0.01 “**”, and *p ≤* 0.05 “*”.

**FIG. D.21:**
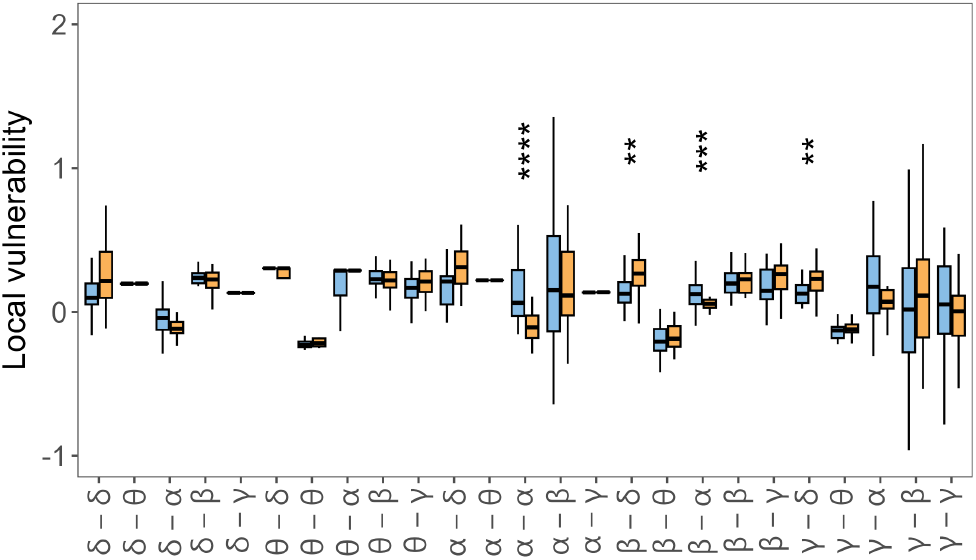
Weighted local vulnerability of HC (blue) and AD (orange) in epoch 3. Significant differences (*p ≤* 0.05) observed in at least ten thresholded networks are encoded by asterisks. The number of asterisks corresponds to the p-value (FDR corrected), i.e. *p ≤* 0.0001 “****”, *p ≤* 0.001 “***”, *p ≤* 0.01 “**”, and *p ≤* 0.05 “*”.

